# High-speed whole-brain imaging in *Drosophila*

**DOI:** 10.1101/2025.06.18.660371

**Authors:** Wayan Gauthey, Albert Lin, Osama M. Ahmed, Andrew M. Leifer, Mala Murthy, Stephan Y. Thiberge

## Abstract

Recent advances in brain-wide recordings of small animals such as worms, fish, and flies have revealed complex activity involving large populations of neurons. In the *Drosophila* brain, with about 140,000 neurons, brain-wide recordings have been critical to uncovering widespread sensory and motor activity. However, current limitations in volumetric imaging rates hinder the accurate capture of fast neural dynamics. To improve the speed of volumetric imaging in *Drosophila*, we leverage the recently introduced light beads microscopy (LBM) method. We built a microscope and a LBM module tailored to fly brain experiments and used it to record brain-wide calcium signals in adult behaving flies at either 28 volumes per second or at 60 volumes per second (when selecting the central brain alone). We uncover fast-timescale auditory responses that are missed with standard volumetric imaging. We also demonstrate how temporal super-resolution can be combined with LBM data to uncover responses to single *Drosophila* courtship song pulses. This establishes LBM as a viable tool for capturing whole-brain activity at high spatial and temporal resolution in the fly.

## Introduction

Across model systems, there has been a major push to develop methods that capture large-scale, high-resolution neural activity distributed across the brain (1–8). These methods are capable of recording neural population dynamics simultaneously from distant regions and different cell types. In mice, where the scattering properties of brain tissue have made two-photon microscopy the method of choice (9), instruments such as 2P-RAM (1) combine remote focusing for volumetric imaging and large custom optics to achieve a large field-of-view that covers a significant portion of the cortex. In *C. elegans*, single-photon spinning disk confocal microscopy has been successful in resolving the activity of each of the ~100 neurons present in the animal’s head (10), including during behavior (11, 12). In larval zebrafish, a larger brain with low scattering properties, it has been possible to use single photon light sheet microscopy (13) to capture activity from roughly all 100,000 individual neurons (14), both in immobilized (15) and freely moving (16) preparations. The relatively low temporal resolution of these techniques in zebrafish (~1-4 volumes per second) has not been a major concern, as the current primary limiting factor of these studies has been the slow varying fluorescence signal associated with nuclear-localized calcium sensors.

In flies, single-photon light-sheet approaches such as SCAPE have achieved large-volume, high-speed (10 volumes per second) imaging (7, 17). However, the high scattering properties of the fly brain limit the ability to report neural activity beyond a depth of 70 to 90 µm, capturing only a fraction of the 140,000 neurons in the brain (18). Light-field microscopy, a single-photon method which employs microlens arrays and compu-tationally reconstitutes entire volumes from single camera frames, achieves very high speeds (~ 100 volumes per second), but without achieving cellular resolution (~ 5 × 5 × 20 *µ*m) (19–23). For fly brain imaging, the preferred volumetric method, offering cellular-scale resolution at depth, has been 2-photon scanning mi-croscopy coupled with a fast piezo objective (6, 24). This method is largely insensitive to tissue scattering and achieve subcellular resolution at depth. Recent advances have improved the speed and sophistication of population 2p imaging in *Drosophila* larvae (25). In adult flies, however, the temporal resolution remains low in comparison to neural dynamics, at a few volumes per second when imaging brain-wide.

The recent development of two-photon light-bead microscopy (LBM) (26) represents a major improve-ment over existing approaches to recording from large swaths of the brain at cellular resolution. LBM relies on a customized optical cavity that splits an incoming beam from a low repetition rate pulsed laser to generate a set of several tens of axially separated and temporally distinct foci (also named ’light-beads’) in the sample. LBM does not suffer from the limitations of other high-speed strategies. Methods relying on extended foci (27–30) require signal disambiguation and are limited to sparse activity. Methods relying on multiple foci and spatially resolved detection (31) are limited by scattering-mediated crosstalks. In LBM, the temporally distinct excitation at each foci, several nanoseconds from each other, eliminates both of these limitations. These foci are moved horizontally through the sample using a conventional resonant scanning method, gen-erating multiple imaging planes near-simultaneously. A critical element of the approach is the optimized temporal sequence in which the pulses arrive at each foci. The sequence is designed such that the source of fluorescence can be identified unambiguously on the basis of the timing in which photons arrive. Further, the signal integrated in a voxel is gathered after a single laser pulse during the smallest time interval possible, the fluorescence lifetime of the probe. This single-pulse-per-voxel sampling approach maximizes the signal-to-noise ratio per unit of power delivered to the brain, while also freeing up temporal resources that can be used either to image the largest possible volume, increase the imaging rate, or both.

To date, LBM has only been applied to the mouse cortex. However, LBM has the potential to address the shortcomings of existing volumetric calcium imaging methods used in small animals and, in particular, could have an immediate impact in the case of Drosophila melanogaster which would benefit from improvements in both the speed and the fraction of the brain being imaged. This would also enable imaging of faster ge-netically encoded voltage indicators (32), and would further facilitate relating brain-wide neural dynamics to the recently completed whole-brain connectome for *Drosophila* (18). Finally, even experimental approaches where only a few cell types are labeled would benefit from whole-brain imaging as circuits often comprise neurons with process distributed throughout the brain (33, 34).

LBM was originally developed as an added module to the 2P-RAM microscope, replacing the original microscope remote focusing capability with the LBM optical module. Directly applied to Drosophila brain dimensions, its original configuration, which uses only 120 pixels per line, would result in a pixel size of more than 2.5µm. This would not be adapted to the *Drosophila* brain which requires a different configuration. Two labeling strategies have been used in *Drosophila*. Experimental approaches where GCaMP is localized in the nucleus (7) are not suited to study fast dynamics. In contrast, approaches where GCaMP is localized in the cytoplasm reveal fast dynamics in the neuropils, rich in axons and dendrites. The somas themselves, however, can rarely be exploited as only a minority of them have recordable signals (24) (although see (35)). While one cannot unambiguously distinguish signals from individual neurons, cytoplasmic GCaMP is chosen to reveal activity at all relevant temporal scales. Given this “neuropil-centered” approach, high spatial sampling, on the scale of a micron, is necessary. Therefore, we implement LBM on a different microscope, updating its optics, the scanning system and the optical path between the laser and the microscope to achieve dense spatial sampling of the whole fly brain at high volume rate (See methods for the considerations influencing the choice of laser repetition rate and resonant scanner).

Figure 1 depicts our LBM implementation. We demonstrate whole-brain activity recording at high spatio-temporal resolution (~ 1 × 1 × 10 *µ*m at 28 volumes per second), up to a depth of 300 µm. We employ this microscope to record whole-brain responses to fly courtship song, a highly dynamic stimulus with fast timescales. We uncover widespread responses to auditory stimuli, including neural responses to single pulses of courtship song. In addition, we demonstrate that LBM can be used over tens of minutes of imaging without degradation of animal behavior. Our study demonstrates the application of LBM to volumetric imaging in the *Drosophila* brain, a powerful model for systems neuroscience.

**Figure 1.**
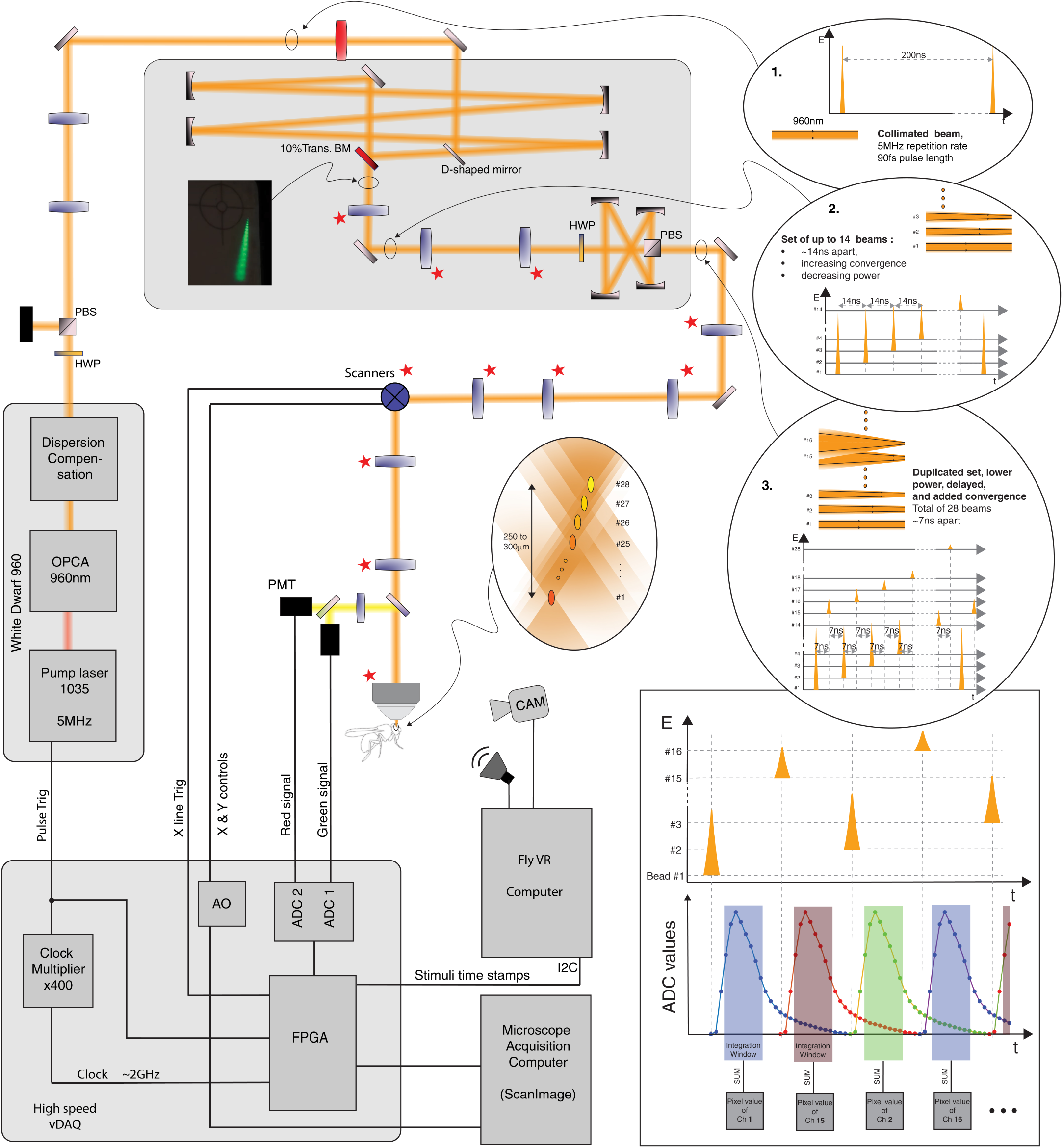
LBM schematics. Our system uses a 5MHz laser which enters the first LBM cavity as a collimated beam (1). A set of 4 concave mirrors and a nominal 10% beam splitter generate a set of 14 beams, 14 ns apart, of increasing convergence and decreasing power (2). A second set of 4 mirrors create a copy that is delayed by 7ns, and where additional convergence is added. The result (3) is 28 beams temporally separated by 7ns of increasing convergence and decreasing power, which when relayed to the sample generate a set of foci (‘light-beads’), 250-300µm long. The deepest foci has higher power, the most shallow one, the lowest power. The temporal separation between ‘beads’ is sufficient to determine the origin of the signal without ambiguity (bottom right panel). Our system includes a computer equipped with a High Speed vDAQ (MBF) and running ScanImage, and a ‘behavior’ computer, generating the audio stimuli and recording the behavior of the animal, as well as time stamps recorded by the ‘Imaging’ computer. The red stars indicate elements that differ from the original LBM implementation from (26). Optical elements are listed in Supplementary Fig.S1

## Results

Light-Bead Microscopy (LBM) (26) was first implemented in the large field-of-view 2P-RAM microscope (1), which uses specialized large optics and is tailored for cortical neural activity fluorescence imaging of the mammalian brain. We adapted the LBM approach for brain-wide imaging at high-spatial sampling in behaving *Drosophila*; our aim was to collect neural activity from the entire brain (using pan-neuronal expression of GCaMP, see Methods) along the medial/lateral and dorsal/ventral axes (spanning roughly 650µm by 250µm, respectively), and as much activity as possible at depth along the anterior/posterior axis (spanning roughly 250µm) (Fig.2A-C). Our field of view included the central brain and both optic lobes. We used a more conventional microscope, which, although built in-house, includes only standard components and a common commercial objective for a roughly 1 mm diameter field of view. This necessitated the modification of multiple elements that relay the foci generated in the LBM cavities to the imaging plane of the microscope.

**Figure 2.**
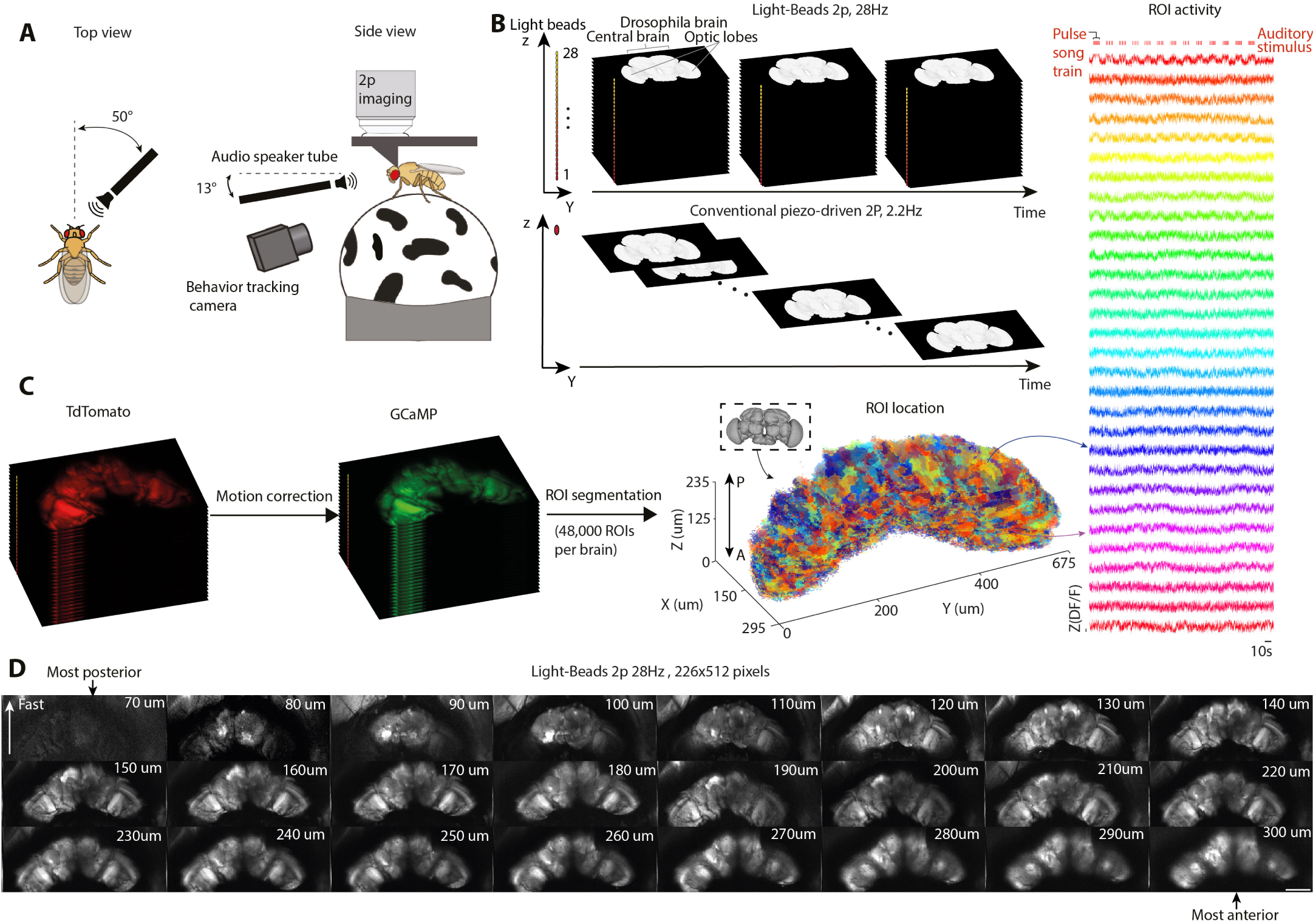
LBM enables whole brain, high frame rate volumetric imaging in *Drosophila*. **A)** Experimental setup. See Methods for more information on sound delivery and preparation of flies for imaging. **B)** Approaches compared in this study for whole-brain imaging in *Drosophila*. In LBM (top) the laser is split into multiple foci, in our case up to 28 (yellow and red dots numbered from 1-28). Practical constraints (see Methods) restrained our use to 24 beams only. These foci, scanned horizontally, probe the individual planes near-simultaneously, thereby imaging the entire sample at once (at a rate of 28 volumes per second). In piezo-driven 2p the single beam scans each plane sequentially (2.2 volumes per second). **C)** Image processing and signal extraction. (left) Images were motion-corrected using the anatomical channel (tdTomato) and the transformation was applied to the activity channel (GCaMP). (middle) A total of 48,000 ROIs were extracted per brain (2000 per plane) (see Methods). (right) Z-scored Δ*F/F* over time for a subset of 30 randomly chosen ROIs from one brain. **D)** Averaged image for each of the 24 planes extracted with LBM for one fly. The number indicates the depth in the sample. The scale bar equals 100µm. The “Fast” arrow indicates the scanning direction of the resonant scanner.

Our LBM implementation, for which the configuration is depicted in Fig 1 and in the Supplements (Text and Fig. S1), results in a 295 µm width by 675 µm length by 235 µm depth volume scanned with high density sampling at a rate of 28 volumes per second. Using a 5 MHz repetition rate laser and an 8 kHz resonant scanner, our system was configured to simultaneously scan up to 28 different planes with 226 pixels per line. When images are acquired using 512 lines, the resulting volumetric imaging rate is 28 volumes per second, with an average pixel size of 1.3 µm, parameters chosen to adequately capture neuropil activity. In our LBM implementation, the number of light-beads was purposefully reduced to 24 because of limitations on available commercial optical beam-splitters (see Methods). The separation between imaging planes is tuned in the first optical cavity of the LBM module, and was set to a value of 9.5 µm. This value matches the average axial resolution of the different foci, which is 9 µm (see Fig. S2). Fig.2D shows the 24 planes imaged simultaneously. In this example, we chose to position the top imaging plane 70 µm below the posterior surface of the brain. This was done to ensure that we captured the most anterior parts of the brain, containing the processes of auditory receptor neurons in the antennal mechanosensory and motor center (AMMC) (18, 24). This is illustrated in Supplementary Fig. S3 and S4, which demonstrate the resolving capabilities of our system, and highlight its ability to capture morphological features, even in the deepest planes in the brain.

Our comparative system, a piezo-driven microscope built according to the current state-of-the-art two-photon approach for *Drosophila* brain-wide imaging, has a high and constant axial resolution (5 µm) (6, 8). About fifty sequentially scanned planes are used to cover the depth of the brain, bringing the volume rate to 2.2 Hz (Fig.2B), more than an order of magnitude slower than LBM.

In what follows, we compare the capabilities of the two volumetric imaging approaches to measure the rapid and widespread neural dynamics associated with auditory stimuli. We imaged head-fixed female flies walking on an air-supported spherical treadmill and presented auditory stimuli in the form of artificial courtship song pulse trains; male flies generate courtship songs comprising two main modes, sine and pulse (36, 37) (Fig.2A). The pulse mode consists of trains of species-specific pulse waveforms separated by species-specific interpulse intervals(38). In *Drosophila melanogaster*, pulse durations are typically 10 ms long, with a peak-to-peak interval of 36 ms.

First, we presented pulse trains at different train repetition frequencies varying from 0.25 to 5 Hz (Fig.3A); each train consisted of pulses separated by the species-typical IPI of 36 ms. We simultaneously recorded calcium activity using either the LBM approach or the conventional piezo-driven two-photon approach (Fig.2B). The subsequent segmentation of regions of interest (ROI) (Fig.2C) is described in the Methods. The imaged flies carried both the calcium sensor GCaMP6f (39) expressed pan-neurally under the nsyb-Gal4 promoter and a secondary fluorophore, used for motion correction (see Methods and Fig.2C).

**Figure 3.**
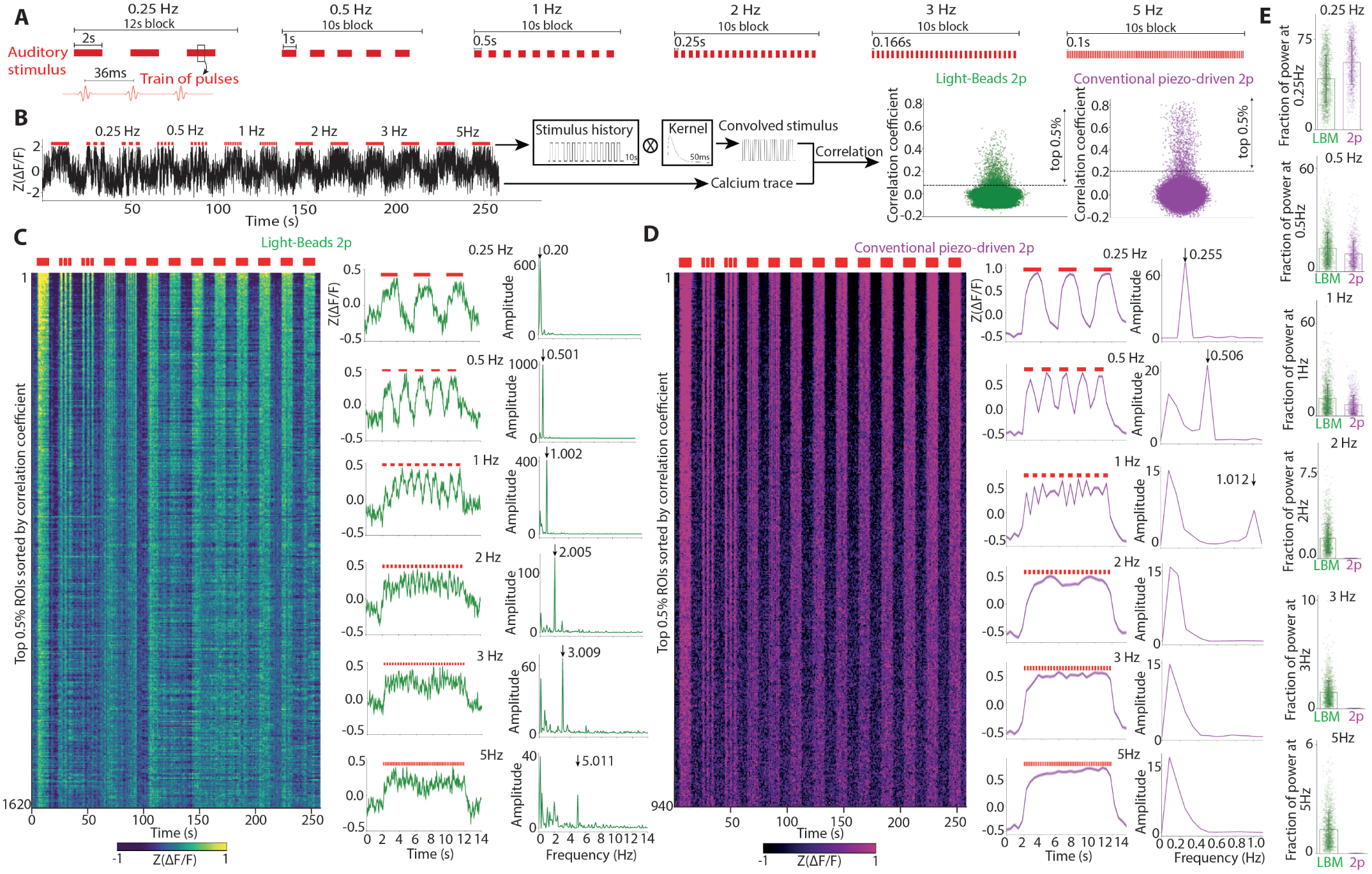
LBM captures fast dynamics across the entire *Drosophila* brain. **A)** Auditory stimulus contains 13 blocks, each 10s long. The first block is a continuous 10s pulse train, with species-typical pulses separated by 36ms IPI (inter-pulse intervals), followed by blocks of pulse trains, with intervals between trains increasing in frequency from 0.25Hz to 5Hz. The duration of each pulse train within a block was adjusted to generate the desired frequency. **B)**.Representative z-scored Δ*F/F* from a single ROI (left) and method for identification of stimulus-modulated ROIs (middle); a binarized version of the stimulus is convolved with a kernel with the temporal properties of GCaMP6f to generate a template. Correlation coefficients between the template and all ROIs are computed. For each imaging modality, the top 0.5% of ROIs with the highest correlation to the convolved stimulus were extracted (right). Dotted line indicates the cutoff. **C)**.(Left) Activity of the top 0.5% of ROIs extracted with LBM and sorted by correlation coefficient (highest on top). (Middle) Z-scored mean activity of the top 0.5% ROIs during each stimulus block of the indicated frequency. Shaded area represents standard error of the mean (SEM). (Right) Fourier spectrum of the z-scored mean activity during each block. Arrow indicates expected peak if activity is stimulus-locked. **D)**. Same as C for conventional piezo-driven 2p. **E)** Bar plot of fraction of power at the indicated stimulus frequency for each imaging modality. A total of 1620 stimulus-locked ROIs were found in the LBM data acquired over 3 animals, each imaged in six 5-minutes long trials. A total of 940 stimulus-locked ROIs were extracted from the piezo-driven 2p data over 2 animals imaged during four 5-minutes long trials.

To extract neural activity correlated with the auditory stimulus, for each ROI, we calculated the correlation coefficient between calcium signal and a filtered auditory stimulus (24)(Fig.3B), and selected the ROIs exceeding a given threshold, which was optimized for each animal (Supplementary Fig.S5B-C and E-F, see Methods). We found that auditory ROIs were widespread, covering the entire brain volume (Supp. Fig.S5A and D), and comprised a diversity of response timescales, consistent with previous work (24).

To determine whether LBM could extract fast timescale activity, we inspected the 0.5% ROIs with the highest correlation to the stimulus (Fig.3B). This subgroup of ROIs is primarily concentrated in the AMMC and wedge (WED) neuropils (Supp Fig.S6); these brain regions receive direct inputs from auditory receptor neurons (Johnston’s Organ neurons) in the antenna and are in the deepest planes we imaged.

Mean activity of ROIs with the highest stimulus-correlated activity extracted from LBM imaging tracked the auditory stimulus at all frequencies presented (Fig.3C). In contrast, the mean activity of the most stimulus-correlated ROIs extracted from piezo-driven 2P imaging failed to time lock to stimuli above 1 Hz (Fig.3D). Fourier analysis of the mean activity confirmed these observations (Fig.3C and D). We repeated these analyses for individual ROIs and found similar results (Fig.3E and Supplementary Fig.S6). The secondary fluorescence (tdTomato) signal was used to verify that stimulus correlations do not appear due to motion artifacts (Supplementary Fig.S7). ROIs extracted from both imaging modalities showed a high fraction of power at frequencies up to 1Hz, while only ROIs extracted with LBM showed power at frequencies above 1Hz (Fig.3E). Our results demonstrate that for brain-wide imaging, LBM can reveal fast neural dynamics in *Drosophila* that standard 2P cannot. We found that stimulus-responsive ROIs are found primarily at depth (e.g., in anterior brain regions like the WED and AMMC) (See Fig.S8).

Next, we investigated whether LBM’s high frame rate could resolve even faster dynamics, namely neural responses to the courtship pulse song inter-pulse interval (IPI). This song feature varies across species (40) and is known to be important for song perception (38, 41). Previous electrophysiological recordings of early auditory neurons in the *Drosophila* brain reveal that some single neurons can track individual song pulses (42). However, whether such activity can be uncovered with calcium sensors is not known. Our stimuli contained the *Drosophila melanogaster* species-specific inter-pulse interval (IPI) of 36 ms (or 27 Hz) (43). To capture neural dynamics at such timescales, we adjusted our LBM’s field of view by reducing the number of lines scanned. This resulted in imaging only the central brain (excluding the optic lobes, but at full depth) at 60 volumes per second. We then recorded neural activity of head-fixed and walking female flies expressing both tdTomato and GCaMP8m (44) pan-neurally while presenting blocks of 2s auditory pulse trains (Fig.4A). We used GCaMP8m for these experiments because it has faster kinetics (~58ms half-rise and ~137ms half-decay times) than GCaMP6f (44). We found that the activity of some ROIs within a single stimulus block showed a peak in the 27-28 Hz range in their Fourier spectrum (Fig.4B-D). We then analyzed all ROIs that ranked in the top 20% for absolute power in the 27–28 Hz range and exhibited a positive correlation with the stimulus (Fig.4E-G). Across all ROIs, we computed the mean Fourier spectrum and the absolute power during stimulus presentation, before or after the stimulus, and during stimulus presentation but shuffling the activity. We found that the mean Fourier spectrum across all ROIs during stimulus presentation showed a peak at the expected frequency of 27Hz, which was absent when activity was shuffled or when there was no stimulus (Fig.4F). Similarly, ROIs had a statistically significant increase in absolute power in the 27-28 Hz range when the stimulus was present (Fig.4G). This suggests that neural responses tracked individual pulses at 27Hz within the stimulus. These ROIs spanned all 24 imaged planes (Fig.4A).

**Figure 4.**
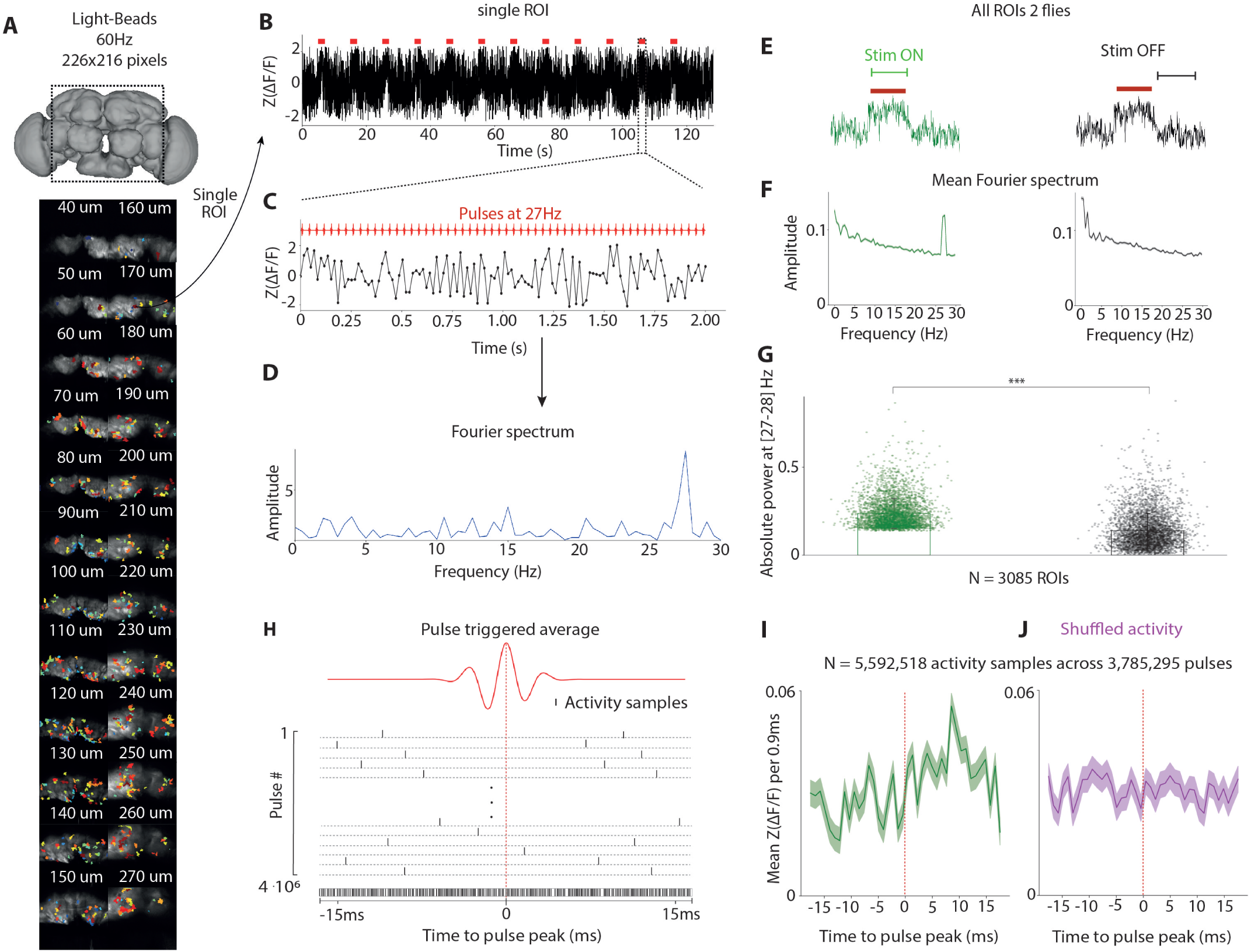
LBM resolves brain-wide neural activity that tracks individual song pulses within pulse train stimuli. **A)** (Top) Field of view (central brain) for the experiment. (Bottom) Maximum intensity time-projections of the 24 planes imaged with LBM at 60Hz overlayed with ROIs that tracks individual song pulse. Number indicates depth in the sample. **B)**. Representative z-scored ΔF/F of a single ROI. **C)** 2s snippet of this ROI during a single block (36ms IPI). **D)** Fourier spectrum of the neural response revealing a peak at 27Hz. **E-G)** Plots of mean Fourier spectrum and absolute power in 27-28Hz in two different conditions: during stimulus presentation (green) and when there was no stimulus (black). Shaded area represents standard error of the mean (SEM). \**p <* 0.05, * * *p <* 0.01, * * \**p <* 0.001. **H)** For each sound pulse (red trace) of each ROIs across all animals, we extracted both the sample time-points within 18ms before and 18ms after the pulse peak, and their fluorescence values. The pulse triggered average was obtained by taking the mean activity per 0.9ms in this temporal superresolution distribution. **I-J)** Pulse triggered average plot of the activity and shuffled activity.The grey area represents SEM. Dotted reed line represents the time of the pulse peak.

While the above analyses suggests that GCaMP8m signals can track individual song pulses, they do not reveal the shape of those responses. To explore this, we calculated a pulse-triggered average by combining and time-aligning all activity samples within ± 18ms of all song pulses across the same subset of ROIs and found that the reconstructed activity peaked at ~9 ms following each pulse (Figs. 4H and I). The pulse-triggered average did not show such a peak when activity was shuffled. In principle, reconstructing the pulse triggered average is possible even with the lower frame rate of conventional 2P (2.5Hz) using a previously published method (45), by leveraging the repeated nature of the stimuli. However, our imaging system requires 30 times less data to generate that same result. Our results highlight that we can not only track fast neuronal activity with LBM, but that by fine-tuning the field of view, LBM speed can be adjusted to experimental needs.

Finally, because we imaged neural activity while female flies walked on an air-supported treadmill (Fig.2A), we could assess how LBM affected fly behavior. In the conventional piezo-driven 2P imaging modality, the animal is exposed to a single focused beam typically between 1 to 15mW in power depending on the depth in the tissue. With LBM, higher overall power is used (about 47mW in our experiments), but unlike conventional 2P, this total power is distributed over multiple foci at different locations, resulting in only moderate photobleaching over long exposures (Supplementary Fig.S9). Despite higher average power, LBM did not negatively impact fly behavior, over 40 minutes of imaging (Fig.S10). For each trial (for either the LBM or conventional 2P), the walking speed was recorded. We found that the walking speed remained consistent across time (Fig.S10B). Notably, there were no significant differences in this metric between the first and last trial, even over 40 minute imaging sessions.

## Discussion

The LBM method, with its reliance on time-multiplexed two-photon foci, is capable of high rate volumetric imaging. With the proper combination of laser, scanners, and optical components, we have demonstrated it can capture activity simultaneously throughout the entire fly brain.

LBM speed is fundamentally limited only by the fluorescence lifetime of the probe, which means that information cannot be extracted from the specimen at a faster rate. However, different LBM configurations can favor different applications. Of particular importance is the choice of resonant scanner frequency and laser repetition rate, parameters which, respectively, fix the number of pixels per line, and the number of light-beads. The original LBM implementation (26) which used only 120 pixels per line would not apply well to the *Drosophila* brain where a high spatial sampling on the scale of a micron is necessary to properly capture the activity from the entangled neural processes, or neuropil, as well as individual somas. Using the Demas et al. configuration would result in sampling the brain dorsal/ventral axis (for the positioning we used, see Methods) only every few microns. To unlock the power of the light-beads for the fly, we have used a configuration of resonant scanner frequency, laser repetition rate and optical elements that together achieve high-density sampling in the imaging planes (1.3um), a separation between planes commensurate with the axial resolution (9um, Fig.S2A), and a high temporal resolution (tens of milliseconds). In addition, properly shaping the beams to optimize the resolution and the separation between planes required modifying the optical path leading to the microscope.

In the future, several refinements of the microscope could further improve our ability to extract neuronal dynamics. These include increasing the signal-to-noise ratio (SNR) of the images and improving control of the light-beads power distribution. With the power used in our experiments, images acquired are typically in the single-photon per pixel regime: most pixels result in zero to only a few photons being detected. Future brighter probes will, of course, be beneficial. Fast detection systems with lower noise, such as Hybrid Photo-Detectors (HPD) which differentiate events with zero, one, and multiple photons better than photomultiplier tubes (PMTs) should in principle increase the SNR. Their higher speed will also help reduce further the low cross-talk (Fig S2C). The power distribution of the light-beads is inherited from the fixed-transmission coef-ficient of the beam splitter present in the cavity, which is nominally 10% in our case. The first light-bead, the deepest in the brain, has the highest power, and each following bead has ~ 10% less power than the previous. This power profile aims to compensate for tissue scattering and generate imaging planes of similar brightness. This was largely achieved, although overall brightness differences remain (Supplementary Fig.S11). No alter-native commercial beam-splitter, in particular of lower value, could be identified to test further improvements in brightness homogeneity across planes. Future implementations of LBM could benefit from the design of custom beam splitters.

We have shown that our LBM implementation enables volumetric imaging of the entire *Drosophila* brain at 28 volumes per second. This rate can be increased to 60 volumes per second by restricting the field of view to just the central brain, and could be increased further with smaller fields of view. Animal behaviors and sensory processing often occur on millisecond timescales. We demonstrated that the fly brain can reliably encode high-frequency auditory stimuli and resolve single song pulses that occur in natural song at a rate of 27Hz. Such observations demonstrate a major capability of LBM over previous imaging methods. Yet, faster dynamics do exist in the *Drosophila* brain, for example in the visual system (46) or olfactory system (47, 48). Moreover, the spiking activity of individual neurons can be recorded with promising new voltage sensors compatible with two-photon imaging (32). Recording such activity requires fast scan rates in the hundreds of Hertz. It is possible to adapt LBM to voltage indicator applications, but at the expense of either restricting the field of view, or reducing the density with which the sample is being probed. The latter could be done, for example, using different scanners, or potentially, by adopting alternative scanning patterns (25). Note also that LBM is compatible with optogenetic perturbation methods. The combination of both could in principle allow monitoring the effect of stimulation or inhibition of single neurons on the Ca dynamics across the brain.

The ability to extract neural activity across the entire brain with high spatial and temporal resolution represents a significant technological advance that opens new avenues for studying how neural circuits process fast and complex stimuli or generate behavior (49). LBM has now been applied successfully to mice(26) and *Drosophila* (this study), and studies of other animal models would likely benefit from the technique as well. In particular, LBM represents an alternative to light-sheet imaging, improving the temporal resolution as well as the imaging depth. While we applied LBM to imaging brains with pan-neuronal GCaMP expression, we could have alternatively used available genetic driver lines in *Drosophila* (50) to obtain sparser GCaMP labeling, and assign activity to specific cell types. This would enable making connections with the recently completed connectome of *Drosophila* (18). The connectome reveals potential paths through which information may flow, but functional imaging data are needed to determine how signal propagation occurs. Because LBM collects multi-scale information about brain dynamics, from the micron scale to the whole brain, at temporal resolution sufficient to extract most dynamical events, it can be used to uncover the principles relating the cellular diversity of brains to their function.

## Methods

### LB microscope design considerations

In LBM, a single laser pulse is used per pixel. In these conditions, the “width” of a pixel becomes the lateral distance traveled by a beam between 2 successive pulses.

For the application to the fly brain, we argue that it is best that the distance covered between 2 successive pulses be close to the width of the PSF (~ 1µm). The fly brain is for most part a dense web of submicron processes. The volume of a 2P-PSF already can intercept multiple processes. Moving the beam less than the width of the PSF between successive pulses would imply probing the same processes multiple times. Moving it more than the width of the PSF would result in some processes not being imaged at all. The 2P-PSF width being typically on the order of a micron, the “pixel” size should be set preferably to that same amount. And since the width of the fly brain is on the scale of ~ 250µm, a combination of laser repetition rate and scanning system resulting in 200 to 250pixels per line would be ideal. The same argument can be made regarding the axial direction. 2P-PSF are elongated along the optical axis direction. If the separation between planes is set on the order of the length of the axial PSF, then all thin processes of the brain can be maximally captured. The brain is ~ 250 − 300µm thick in the orientation it is being imaged.

Multiple combinations of laser repetition rate (Fl) and resonant scanner frequency (Fres) can achieve that. The number of pixels per line (Np) is equal to

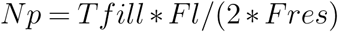

where Tfill is the temporal fill fraction of the image, usually 0.7. However the choice of the laser repetition rate affects also directly the maximum number of light-beads (Nlb), or planes, that can be configured in the system without encountering signal source ambiguities. That maximum number is

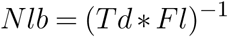

where Td is the decay time of the fluorescence signal, which includes the cumulative effects of the lifetime of the fluorescent probe and the dynamics of the detection system. In the case of GCaMP, Td is around 7ns (26). Finally, these parameters also influence the imaging rate. In LBM, the volume rate and frame rate are the same, and are determined by the resonant scanner frequency and the adjustable number of lines.

Only 3 types of resonant scanners are commercially available: 4, 8 and 12kHz. And laser repetition rates come only in limited possibilities, typically 1, 4, 5, and 10MHz. The combination of a 8kHz resonant scanner (62.5us per line) and a 5MHz laser (200ns) results in about 220px per line and a maximum of 28 light-beads, which if equally distributed across the depth of the animal’s brain results in a 8-10µm separation. To cover the long axis (~ 680µm) of the brain with the same density as the short dimension, about ~500 lines are necessary, resulting in a 30Hz volume rate. Using a 4kHz resonant scanner with a 4MHz laser repetition rate would result in an alternative configuration with 355 pixels per line (more than necessary), 35 planes 7µm apart, for a volume rate of ~15Hz. Finally, a 12kHz resonant scanner (as in the original LBM configuration) combined with a 10MHz laser would lead to 295 pixels per line and only 14 light-beads, separated by 15um, and a volume rate of ~45Hz. Of the three configurations, the first two seem the most adapted, with an advantage for the first being its higher speed.

### Experimental animals

3-6 day old female *Drosophila melanogaster* with genotype *norp36A/+; UAS-GCaMP6f, UAS-myr::tdTomato/+; nsyb-gal4/+*, which express both GCaMP6f and tdTomato pan-neuronally, were used for the LBM experiments at 28 Hz and conventional 2P imaging experiments. Flies with genotype *+/+; UAS GCaMP8m/+; nsyb-gal4/+*, which express GCaMP8m pan-neuronally, were used for the 60 Hz imaging experiments.

### Animal preparation

Female flies were chilled on ice, then placed in a Peltier-cooled “sarcophagus” held at 4°C, with the head of the animal placed in a 3D printed holder. The head of the fly was tilted at a 80° angle relative to the thorax and restrained with UV-cured glue. The arista were exposed so that the flies were able to hear. The holder was then filled with saline (103 mM NaCl, 3 mM KCl, 5 mMTES, 1 mM NaH_2_PO_4_, 4mM MgCl_2_, 1.5 mM CaCl_2_, 10 mM trehalose, 10 mM glucose, 9 mM sucrose, and 26 mM NaHCO_3_) and the cuticle on the back of the head was removed using a sharp needle and fine forceps (Dumont 5SF). Fat and trachea were removed before imaging the brain.

### Animal auditory stimulation and behavior recording

Auditory stimuli were delivered in a localized manner using a headphone speaker and sound tube, as previously described (24, 42). In these experiments, the sound tube was positioned in front and to the right of the fly, at a distance of 2 mm from the head and an angle of 50 degrees. Audio stimuli were generated on a dedicated computer using custom MATLAB code and presented to the animal at a rate of 44100 samples/s using the flyVR software package (github.com/murthylab/fly-vr), which synchronizes stimulus delivery with behavioral quantification. The fly was positioned under the microscope on an air-suspended 9 mm diameter foam ball, which was illuminated with IR LEDs and tracked using an IR camera acquiring at 100 Hz. The animal’s locomotion was extracted using Fictrac (51). This behavioral setup was the same for both LBM or piezo driven 2p.

To synchronize the imaging computer with the stimulus and behavior computer, we employed an Inter-Integrated Circuit (I2C) controlled by a NI USB-8452 interface. At a rate of 100 Hz, a digital signal containing the Fictrac frame number was sent from the stimulus and behavior computer to the imaging computer. This metadata is directly embedded in the image files generated by Scanimage during imaging.

### LB microscope assembly and imaging

We modified the optical assembly design described in Demas et al.(26), employing a different set of scanners and optical elements better suited for imaging fly brain volumes (Fig.1 and Supplementary Fig.S1).

The axial multiplexing module, which splits the laser beam into multiple beams, is composed of two ring cavities. The first optical cavity includes four concave mirrors (500 mm focal length) positioned such that upon each circulation through the cavity the beam accumulates additional convergence. A partial beam splitter placed in the beam path allows a fraction ( 9%) of the beam to escape the cavity every time it circulates, generating a series of spatially separated and temporally offset beams. This set of beams is then sent through a half-wave plate and a polarizing beam-splitter cube. A high portion of the power is transmitted through the cube and sent toward the microscope objective where it forms the first and deepest set of light beads. The portion reflected by the polarizing beam-splitter cube is traveling through the second optical cavity, where a large amount of convergence is added and the pulses are delayed by another ~ 7 ns. This second set of beams is returned to the microscope path to form the second set of light beads, located directly above the first one, and therefore probing the most shallow layers of the tissue.

The separation between imaging planes (*z*) is affected by two parameters: the added path length in the first cavity (*Z*) and the overall magnification (*M*) between the first cavity and the sample. These parameters are related by *Z* = *z* ∗ *M* ^2^. Our microscope employed a 25x water immersion objective (Olympus XLPLAN-N-25X) with a focal length of 7.2mm, a 375mm tube lens, and a 100mm scan lens. The objective back aperture was conjugated to 5mm clear aperture scanners. We combined these components with intermediate optics to achieve an overall magnification of *M* = 156. We tuned the cavity added path length until we achieve a z-separation of *z* = 10 µm between planes. The estimated added path length in the cavity is about *Z* ~ 240 mm. Beam expansion through the system is x1.66 before reaching the scan mirrors, resulting in an XY-resolution of 1µm (See Fig.S2A). The axial resolution varies from 5µm for the deepest planes to 14µm for the shallowest, averaging 9µm.

LBM requires a laser with a lower repetition rate than that used in a conventional two-photon microscope. We chose a 960nm laser (White Dwarf, Class5) operating at a repetition rate of 5MHz. As demonstrated in Demas et al (26), a separation of ~ 7ns between pulses is sufficient to properly identify the fluorescence signal source among the different beads. Applying the same strategy, up to 28 different light beads can be generated simultaneously in our system. Coupling this laser with an 8 kHz resonant galvo scanner (CRS8, Cambridge Technologies), we can achieve a maximum of 226 pixels per scan line. Recording images with 512 lines results in a volumetric rate of ~ 28 Hz. Reducing the number of scan lines gives us a proportionally faster frame rate.

By orienting the animal’s body axis parallel to the resonant scanner axis, the 226 by 512 pixels image format matches the aspect ratio of the fly brain. Consequently, a high spatial density sampling is obtained in the two directions of the planes with an average pixel size of 1.3µm on both axes.

The microscope is run using ScanImage 2022 (MBFbioscience). Our acquisition system includes a High-Speed vDAQ (MBFbioscience), digitalizing the signal at a rate of 2 GHz. This high sampling rate allow to differentiate signal coming from one pulse (one plane) from the signal of the next pulse 7 ns later (next plane). However, given temporal decay of fluorescent signals is exponential, some amount of cross-talk between planes is unavoidable. Fortunately, it can be reduced by integrating only the sampling time points which coincide with the peak response to each laser pulse. The widths and location of each of the 28 signal integration windows were set to capture these peaks. The remaining cross-talk (see Fig S2C) was comparable to that reported in (26).

The microscope included a laser reflecting dichroic (cutoff 765nm, Alluxa), to separate excitation and detection paths. The detection path consisted of a first filter removing the spurious near-infrared laser light (FF01-720/SP, Semrock) followed by a beamsplitter (FF555-Di03, Semrock) separating green and red fluorescent light further individually filtered (green: FF02-525/40, Red: FF01-593/40, Semrock). Photons were detected with two GAsP photomultiplier tubes (H10770PA-40, Hamamatsu).

### Piezo-driven 2-photon imaging

For our comparisons with conventional piezo-driven 2-photon imaging, we carried out experiments on a custom-built scanning microscope equipped a 8kHz resonant scanner, a 80MHz repetition rate Ti:sapphire laser (Chameleon Ultra II - Coherent) set to 920 nm and a 20x water immersion objective (Leica 20x HCX APO 1.0 NA) mounted on a 400µmtravel piezo objective-mount (Physik Instrumente). The microscope is run using ScanImage 2022 (MBFbioscience). Dissected flies were placed below the objective and perfused with saline. The microscope included a laser transmitting dichroic (FF665-DI02 Multiphoton shortpass Dichroic, Semrock). The detection path consisted of a near-infrared filter (FF01-680/SP, Semrock), a 562 nm beamsplitter (FF562-DI03, Semrock) which separated green and red channels, with a 525/45nm filter (FF01-525/45, Semrock) into the green channel path and a 600/52nm filter (FF01-600/52, Semrock) in the red channel. Photons were detected with two GAsP photomultiplier tubes (Hamamatsu H10770PA-40-S1). For functional imaging, we acquired whole-brain volumes with 256 x 128 x 50 XYZ voxels of size 2.4 x 2.4 x 5 µm, and a volumetric rate of 2.2 Hz. We also employed this microscope to capture high-resolution anatomical stacks of all flies after functional imaging conducted either by LB or piezo-driven 2P recordings. We recorded a total of 50 whole-brain volumes 1024 x 512 x 150 XYZ voxels of size 0.6 x 0.6 x 2 µm.

### Image processing and signal extraction

For both 28 Hz LBM and piezo-driven 2P functional recording, each frame of the anatomical (red) channel was motion-corrected by using ANTS(52) to perform a nonlinear alignment to a temporally averaged red channel image of the brain. The transformations made to the red channel were then applied to the activity (green) channel. For the 60 Hz LB data, no anatomical channel was available. Instead, we used a 6-frame rolling average of the green channel as a proxy for the anatomical channel for the purposes of motion correction. After motion-correction, each voxel was independently bleach corrected and high-pass filtered. Voxels were then aggregated into ROIs (~60 voxels on average) to improve SNR. We masked each z-slice to restrict ROIs extraction to the brain. Then, we employed Ward linkage agglomerative clustering with a contiguity constraint on the z-scored voxels(6), extracting 2000 (LBM) or 1000 (piezo-driven 2p) ROIs for each slice. This segmentation process is fully automated, and its output is manually validated to ensure quality. It is therefore suited for large-scale, high-throughput analysis. To compute the signal from each region, we return to the bleach-corrected volume to extract Δ*F/F*_0_.

## Data analysis

### Identification of stimulus-modulated ROIs

48,000 (LB) and 47,000 (piezo-driven 2p) ROIs were extracted in each trial of each animal, then z-scored to account for differences in baseline fluorescence across animals. To identify stimulus-correlated ROIs, we first generated a target template by binarizing the stimulus sequence and convolving it with a kernel with the temporal properties of GCaMP6f (50ms rise time and 140ms decay time(53)). We then computed the correlation coefficient between the target template and all ROIs. in Fig.3, we extracted the top 0.5% most correlated ROIs. To extract the auditory modulated ROIs shown in Fig.S5, a distribution of correlation coefficients was constructed for each ROI by shuffling the stimulus 10,000 times and computing the correlation coefficient between each shuffle and each ROI. A p value was then computed for each ROI, and ROIs whose p value <0.05 were selected.This was done separately for each animal. The pulse-locked ROIs presented in Fig.4 were extracted by calculating (per ROI) the absolute power in the 27-28Hz range. ROIs were then sorted based on their absolute power in that range and the top 20% were extracted. Out of this subset, only the ROIs with positive correlation with the auditory stimulus were kept.

### Fourier analyses

Discrete Fourier transforms were computed on mean-subtracted activity data using scipy’s ‘fft’ function based on the fast Fourier transform algorithm (54). In Fig.3, Fourier transforms were computed separately on the mean activity of each stimulus block. In Fig.S6 and Fig.4F, Fourier transforms were computed on each stimulus block for each ROI. In Fig.4F, the Fourier transform was computed for each individual ROI, yielding one spectrum per ROI. These spectra were then averaged across ROIs to return a mean Fourier spectrum. For the ”Stim off” condition the Fourier spectrum was computed on a window with same length of a stimulus block centered between two stimulus presentations. In Fig.S6 peak detection was computed on the Fourier spectrum of each block of each ROI. In Fig.3, the fraction of power for each block was calculated by taking the absolute power within ±0.1Hz of the corresponding frequency divided by the total power.

### Pulse-triggered average

To reconstruct the pulse triggered average, we identified all time points within a ±18 ms window around each pulse and extracted the corresponding fluorescence values in auditory ROIs, building a distribution of activity around each pulse. The pulse triggered average was plotted by taking the average fluorescence value in each 0.9 ms bin. This approach was based on (45). For these analyses, the integration time of each individual ROI within a frame was calculated and taken into account. In the “shuffled” condition, the activity within each individual stimulus block was shuffled before reconstructing the pulse triggered average.

### Significance tests

In Fig.4G Welch’s two-sided t-test was employed. Multiple hypothesis correction (Benjamini-Hochberg procedure) was applied to the p-values.

## Data Availability

The raw data for a representative trial can be found at https://doi.org/10.5281/zenodo.17613016 and the pre-processed data can be found at https://doi.org/10.5281/zenodo.17618684. Both can also be found at https://doi.org/10.34770/s5hx-1x75.

## Code Availability

The auditory stimulus was presented using the FlyVR software package, available on Github: github.com/murthylab/fly-vr. Image analysis, signal extraction, and neural data processing were performed using custom Python code available at https://github.com/murthylab/lightbead-analysis

## Acknowledgements

We thank Alipasha Vaziri for advice during the construction of the LBM module and the choice of an appropriate DAQ system. We thank David Tank for his advice on construction of the LBM, and the members of the Murthy and Leifer labs for advice on the project and comments on the manuscript. We also thank Manuel Schottdorf for assistance with digital synchronization, as well as David Turner, David Deutsch and John Stowers (LoopBio) for the development of FlyVR. AL was partially supported by the NSF through the Center for the Physics of Biological Function (PHY-1734030). AML and MM are funded by the Keck Foundation, and experiments in the MM lab were additionally supported by the following grants: NIH NINDS R35, Simons Foundation SCGB award, and NIH BRAIN (R01 NS110060). The project was also funded by the Princeton University Dean of Research Transformative Equipment Initiative.

## AUTHOR CONTRIBUTIONS

WG: Stimulus presentation software, imaging experiments, data analysis, writing

AL: Microscope (piezo-2P) design & construction, stimulus presentation hardware & software, imaging experiments, data analysis, writing

OA: Data analysis

AML: Funding acquisition, project supervision, writing, review & editing

MM: Funding acquisition, experimental guidance, project supervision, writing, review & editing

ST: Microscope funding acquisition, design, construction & validation, writing, experiments

## Supplementary text and figures

### Microscope, Light-Bead module and optical path

#### Laser, repetition rate and number of planes

Our laser system is comprised of a femtosecond amplifier pump laser (Monaco, Coherent, 60 W average power, 5.0 MHz), followed by an optical parametric chirped-pulse amplifier (OPCPA, White Dwarf dual, Class5 Photonics) operating at a wavelength of 960 nm with <100 fs duration pulses up to 0.8 µJ in energy. A pulse pre-compensation unit was integrated within the laser enclosure to maintain a short pulse duration at the sample. It uses two pairs of chirped mirrors (Class5 Photonics) generating an approximate total of 17, 000*fs*^2^ of anomalous group delay dispersion to counteract the calculated dispersion generated by our optics in the light-bead module, the microscope, and the relay optics. A half-wave plate and a polarizing beamsplitter cube (Thorlabs) were used to adjust the laser intensity delivered into the first cavity of the light-beads module.

The laser system could be configured in factory for different repetition rates, and we selected 5MHz (or 200ns period), which allows for up to 28 different beams separated by 7ns. This time interval between the arrivals of each beam pulse is sufficient in the case of GCaMP for negligible cross-talks between planes (see below).

#### Resonant scanner and number of pixels per line

We choose a resonant scanner at 8kHz (CRS8, Novanta). Using bidirectional scanning, the line scan is 62.5 µs long. During this interval, about 312 individual laser pulses arrive in each plane. The typical temporal fill fraction of resonant scanning, meaning the portion of the scanner period that is used to form the image, is about 70%, therefore only about 220 pulses per line scan are used to form the image. At the regime of 1 pulse per pixel, the choice of a 8kHz resonant scanner translates to fixing the maximum number of pixels per line to approximately 220. As detailed in the text, this value matches well the requirement for sampling the tissue at spatial intervals equal or below the lateral extend of the point spread function. Fulfilling this condition minimizes losses in spatial intervals being probed and helps capturing accurately the *Drosophila* neuropil calcium activity.

#### Microscope Excitation Path

The microscope was custom designed using Zemax OpticStudio (Ansys). The scan and tube lenses of the microscope are respectively a 100 mm telecentric f-θ lens (4401-464-000, Linos) and a pair of 750mm achromat (PAC097, Newport) forming an effective lens 375 mm in focal length. The scan and tube lenses were in a 4f arrangement. The objective used is a 7.2mm focal length water immersion lens (Olympus, XLPLAN-N-25X), also in a 4f arrangement with the tube lens. The scan head is positioned such that the resonant scanner is relayed to the back focal plane of the objective by the scan and tube lens system. The field curvature of this microscope is negligible thanks to the 4f design and the use of a telecentic f-theta scanning lens and conjugated scanners.

#### The scan box

This is a particularity of our microscope that has no critical influence on the success of this method, but is reported here for completeness. The scan box we have used could have been substituted for a regular scan box using a 8kHz resonant scanner and a 6mm galvanometer without adverse effect. The scan head we use included three mirrors: two scanning Y galvanometers, with respectively 6 and 15mm clear apertures (6215, Novanta) followed by one resonant X scanner (CRS8, Novanta), a configuration referred to by some microscope manufacturers (e.g. Leica) as X2Y. The additional Y mirror virtually conjugate the X and Y scanning axes. In our custom scan head, the geometrical arrangement of the mirrors is such that conjugation is achieved when the second Y mirror moves an angle twice as big as the first Y mirror. Maintaining conjugation between the scan mirrors, and between the scanners and the entrance pupil of the objective helps avoiding lateral shift of the wavefront at that pupil which, if large, could introduce optical aberrations, adding to the aberrations caused by the non-planar wavefronts as encountered in the light-bead microscope.

#### Location of the light-beads

The relative foci locations were verified before every imaging session using a glass slide covered with a monolayer of inhomogeneously distributed 0.5 µm diameter fluorescent beads. The slide was mounted on a piezo stage describing a slow rising ramp. Images were recorded as the monolayer crosses the different light-bead plans. The mean intensity of each image in the stack was recorded and the peak of that value was used to determine the axial location of the light-beads. Image cross-correlation was used to measure the lateral shift relative to light-bead #1. Fig S2 shows the result of a typical measurement. Variations in the position of the light-beads from one day to another were present, but they were relatively small, and were principally affecting the lateral shift.

#### Lateral shift of the imaging planes

Inherent to the design of the light-bead cavity is the necessity for the center of each beam to be slightly laterally shifted from one another (Fig.S2). This is coming from the necessity to first launch the laser in the cavity and keeping it revolving in the cavity for some number of times, in other words the first pass has to differ from the subsequent ones. In our case, we decided to launch the laser using a D-shaped mirror located close to the focal point of the entry lens (L1). The beam being here very small, i can be reflected very close to the edge of the D-mirror. Subsequent revolutions of the beam pass above the D-mirror (M1) without interactions with it. This cause all beams to be atop of each other in the cavity, separated only by a millimeter or so. This shift relayed to the specimen and de-magnified by a factor 156 (X=x*M) by the optical path elements results in a lateral shift of 6 to 8 microns between successive planes. In our implementation, this shift is almost parallel to the scanning axis of the resonant scanner, the short dimension (226 pixels) of the image.

#### Stability of the beads location

Because of the extended length of the optical path, the stability of the foci location has always been of concern. Room temperature fluctuations, for example, may affect the mirror mounts and subsequently the orientation of the beams reflected. For this reason, we opted for daily calibration of the beam locations using the procedure described above. We found that the relative position of the set of beams generated by the first cavity (beams 1 to 14) is stable and rarely requires realignment. The second set of beams, the one generated by the second cavity tends to drift more rapidly (on the scale of a few weeks). It is usually that whole set that drifts laterally together and eventually requires some repositioning. However, lateral deviations of the positions of individual foci is not particularly critical to the approach as it is always possible to correct for the shift, if needed, during post-processing of the images. More critical though is the relative axial separation between foci. We found this separation to be very stable for the first set of 14 beams and to fluctuate more for the second set, likely because of the added paths and optical elements present on this path.

#### Signal integration windows and Cross-talks

The DAQ samples the signal at 2GHz, consequently, there are 400 sampling points in between 2 successive pulses of the 5MHz laser. Twenty-height virtual channels are programmed in ScanImage with signal integration window width integration windows 4.0ns, 8 successive sample points. The temporal order of the beads laser pulses is bead #1, then bead #15, #2, #16, #3, #17, etc… The fluorescence decay time of the GCaMP and the finite response time of the detector and its downstream electronics is such that the signal reaching the DAQ extends up to 10ns beyond the bead laser pulse, generating potentially some small cross-talk between channels. Because of the temporal order of the beads pulses, Channel 1 signal can contaminate Channel 15 image, which can contaminate Channel 2, etc… The exact crosstalk values from Channels 1 to 14, respectively into Channels 15 to 28 can be measured by blocking all the light traveling in the second cavity, thus ensuring only beams 1 to 14 reach the sample. In Fig.S2 we report the ratio of the image intensities in Ch15:28 to Ch1:14. Imaging flies expressing GCaMP8m, we verified that the average cross-talks in our system is 6.9%, a val;e that can likely be reduced substituting the green PMT by an Hybrid-Detector.

#### Beads Power distribution and available commercial beamsplitters

Ideally, the power within each light-bead is such that similar fluorescence signal intensity is collected on all channels simultaneously. It implies choosing the first cavity beam splitter with a well chosen transmission/reflection split ratio. This coefficient should generate a laser power distribution that exactly compensates for the signal loss resulting from the beads being located deeper in the tissue. The appropriate coefficient depends both on the specific scattering properties of the tissue and the chosen spatial separation between foci. In Demas et al. (26), the authors selected a 10% beam splitter which works well for mice cortex and a separation between plans of about 18µm. We estimate that a transmission coefficient of 5 % would be more adapted to the *Drosophila* brain scattering properties and our target separation of 10µm. Unfortunately, no commercially available beamsplitter with this split ratio were identified. It should be noted that the beamsplitter should also have a low Group Delay Dispersion to avoid laser pulse broadening after the multiple reflections in the cavity, limiting further the choices of commercially available elements. Reducing the incidence angle on the beam splitter can be used to slightly decrease the transmission coefficient. We reached a value of about 7% using that approach.

#### Pulse width

Using a custom-built Michelson interferometer positioned before the first cavity and a diluted fluorescein solution, we measured the laser pulse width of several light beads at the level of the sample. We found the pulses to be between 100 and 120 fs.

#### Image deformation and corrections

When acquiring images with a resonant scanner, and a single laser pulse per pixel as is the case in LBM, the inhomogeneity of the sample probing cannot be easily corrected for. Using a 10µm grid target (R1L3S3P, Thorlabs) and fluorescein solution, we acquired images under the same conditions of scan amplitudes and temporal fill fraction as in our experiments. We measured that pixels in the center of the lines represent 1.61 µm in the sample, and pixels at the start and end of the lines represent 0.65 µm, with an average size of 1.31µm. We corrected for the distortions of the images before performing the calibration of the light-bead positions.

## Image and Data Analysis Supplementary Figures

**Figure S1.**
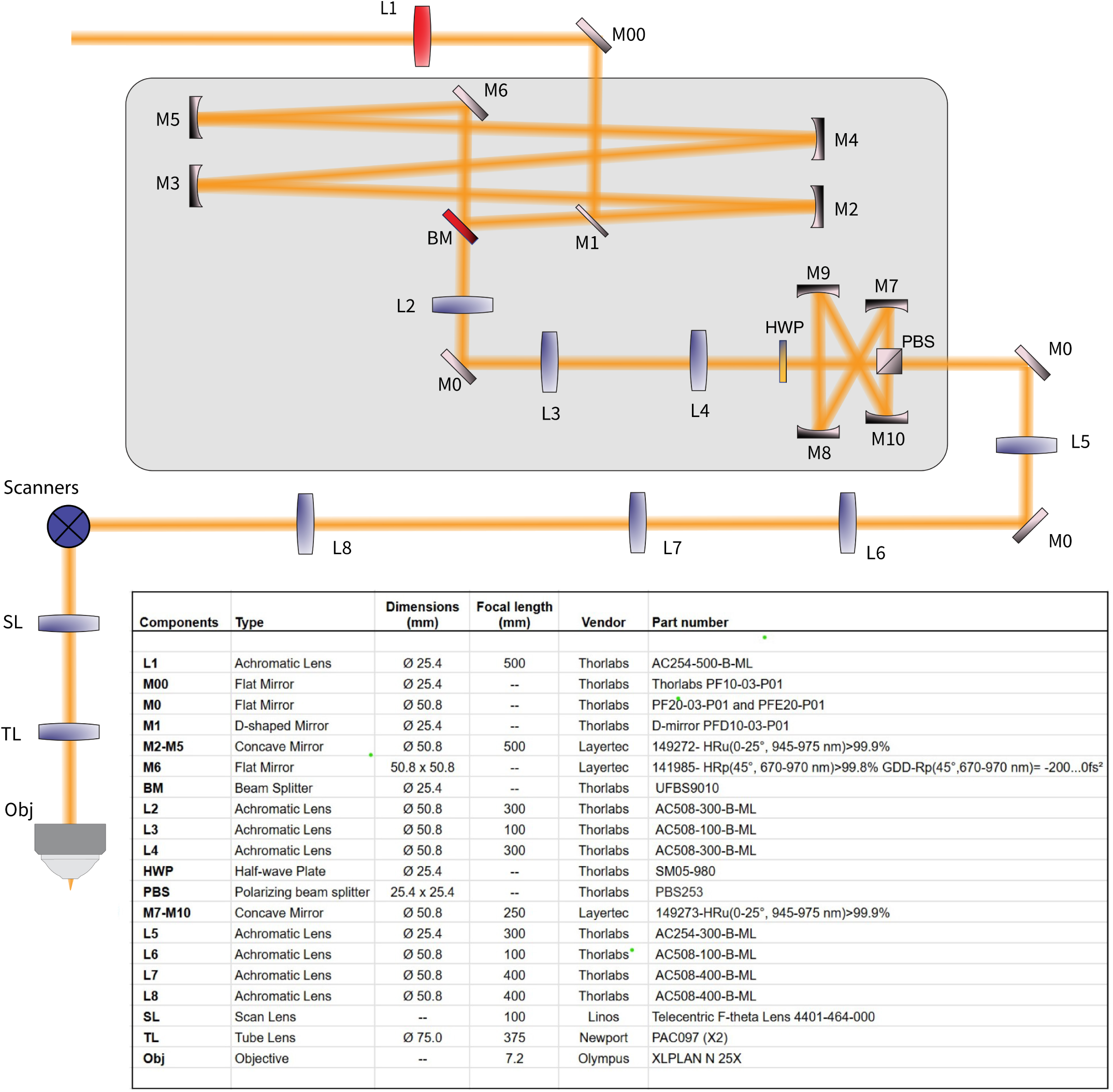
Optical path. Schematic of the optical layout and list of optical elements present in the system.

**Figure S2.**
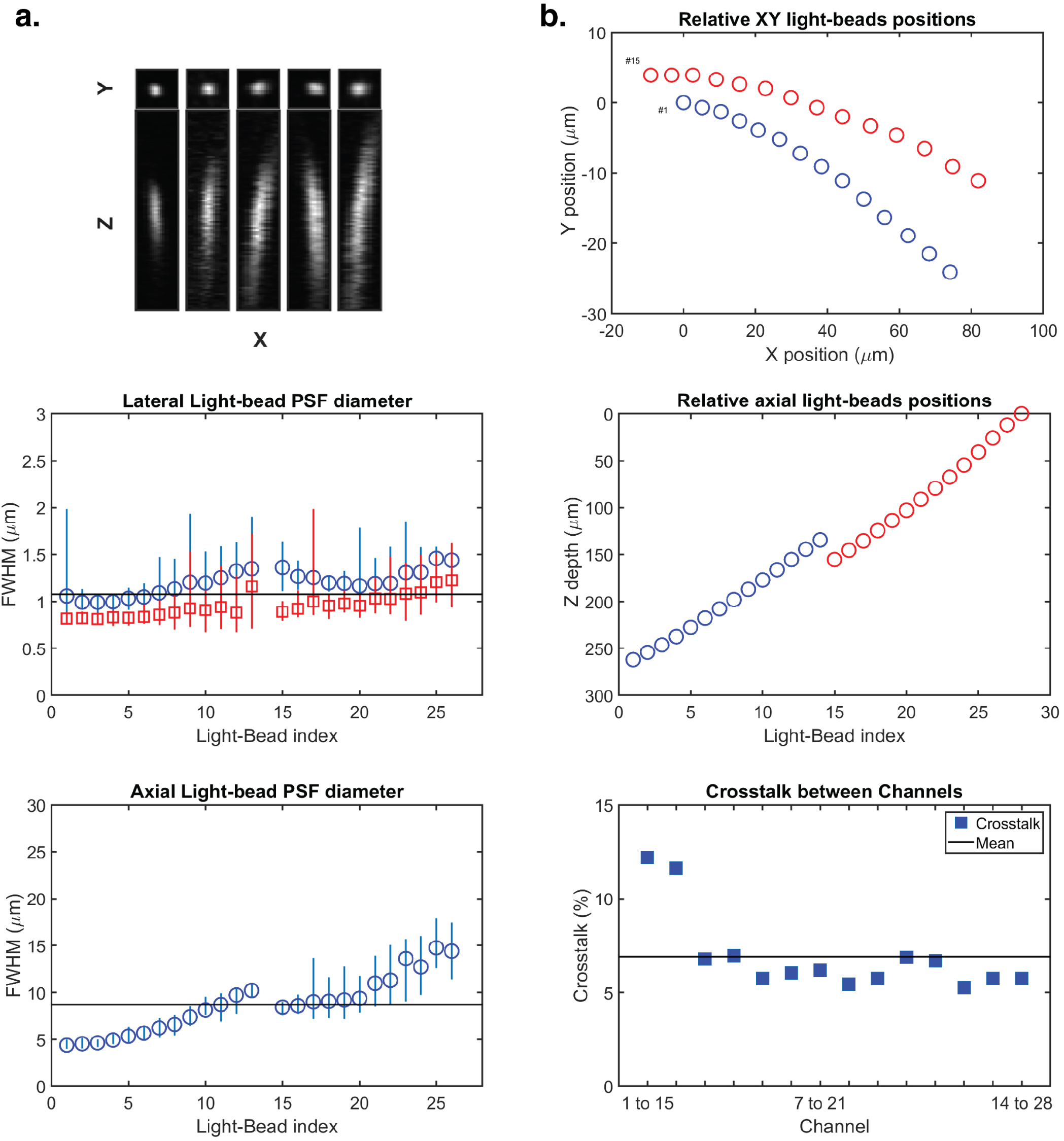
Microscope characterization. **A.** Point Spread Function measurements. Top panels shows 0.5µm green fluorescent beads imaged, from left to right, by light-beads #2, 7, 10, 17 and 24, Mean PSFx (blue circles), PSFy (red squares) (middle panel)) and PSFz (bottom). Bars represent the minimal and maximal values recorded. Lines represent the average value across all ’beads’ **B.** Spatial locations (XY top, Z bottom) of the ’light-beads’ relative to ’light-bead’ #1. Beam #15 was placed at the depth of beam #13, and channels # 13, 14, 27, 28 were later disregarded. **C.** Cross-talks between channels. Line represents the mean value.

**Figure S3.**
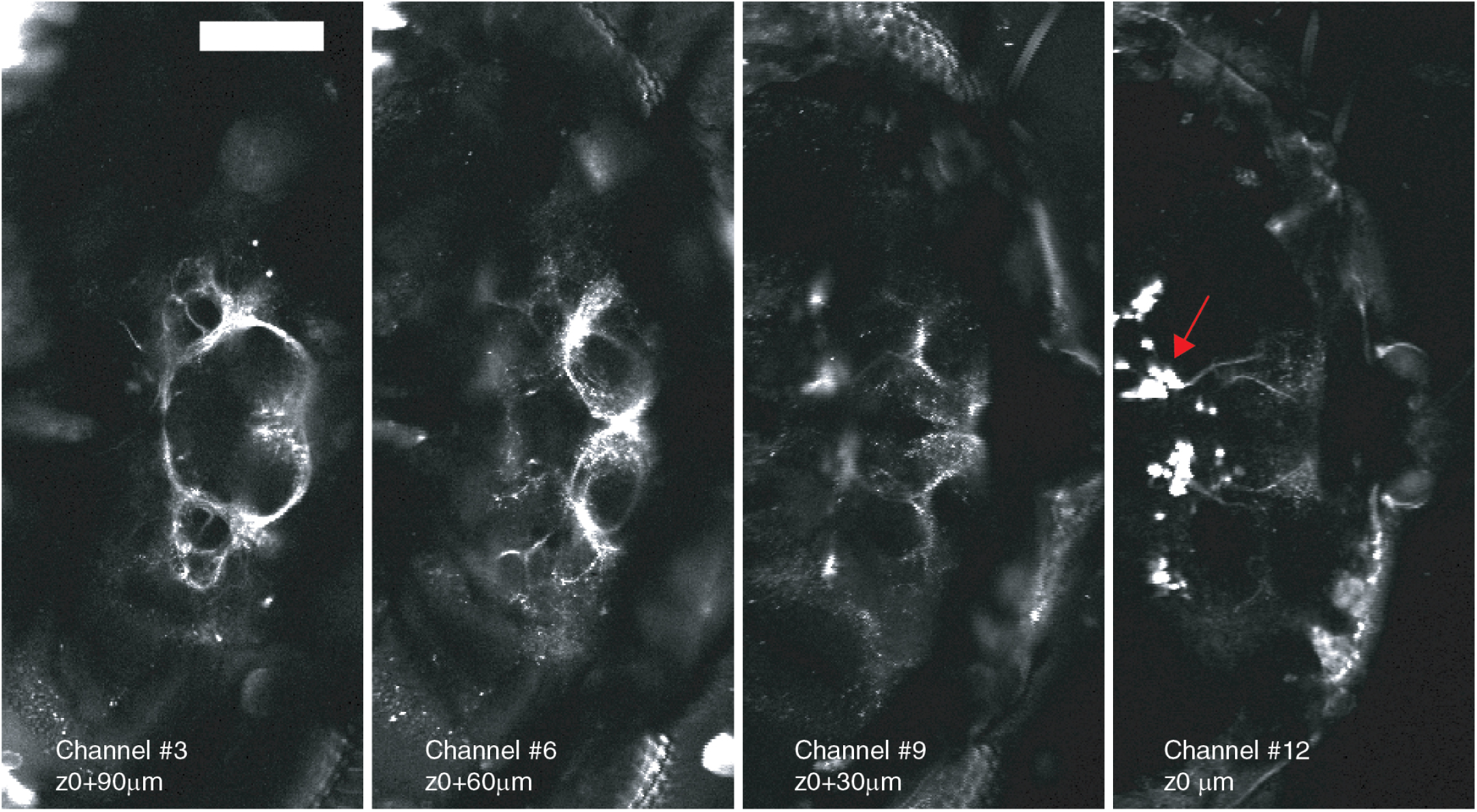
High resolution content. Selected planes showing the result of the sum of hundreds of images from a sparse Gal4 line (Dsx-Gal4 driving GCaMP6m (This driver line labels ~ 100 neurons (55). This highlights the ability of LBM to resolve both cell bodies (red arrow) and processes. Scale bar is 100um.

**Figure S4.**
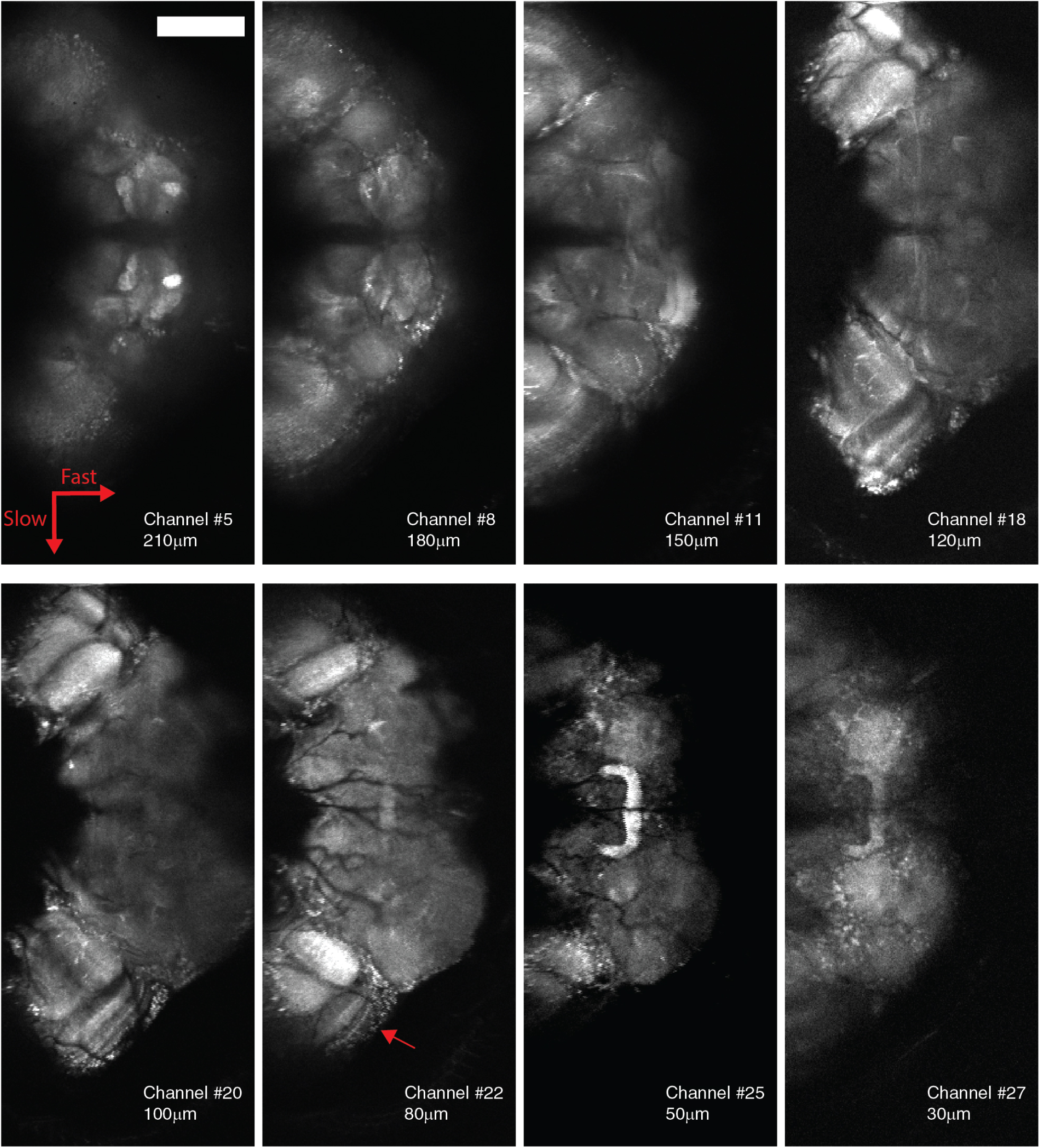
High resolution content. Selected planes showing the result of the sum of hundreds of images from pan-neuronal expression of GCaMP6f. While automatically segmented ROIs likely capture signals from groups of neurons, the spatial resolution of LBM reveals structure within brain regions, such as the layers of the medulla in the optic lobes (thin arrow). Scale bar is 100um. The Fast and Slow arrows indicate respectively the resonant scanner and galvanometer scanning directions.

**Figure S5.**
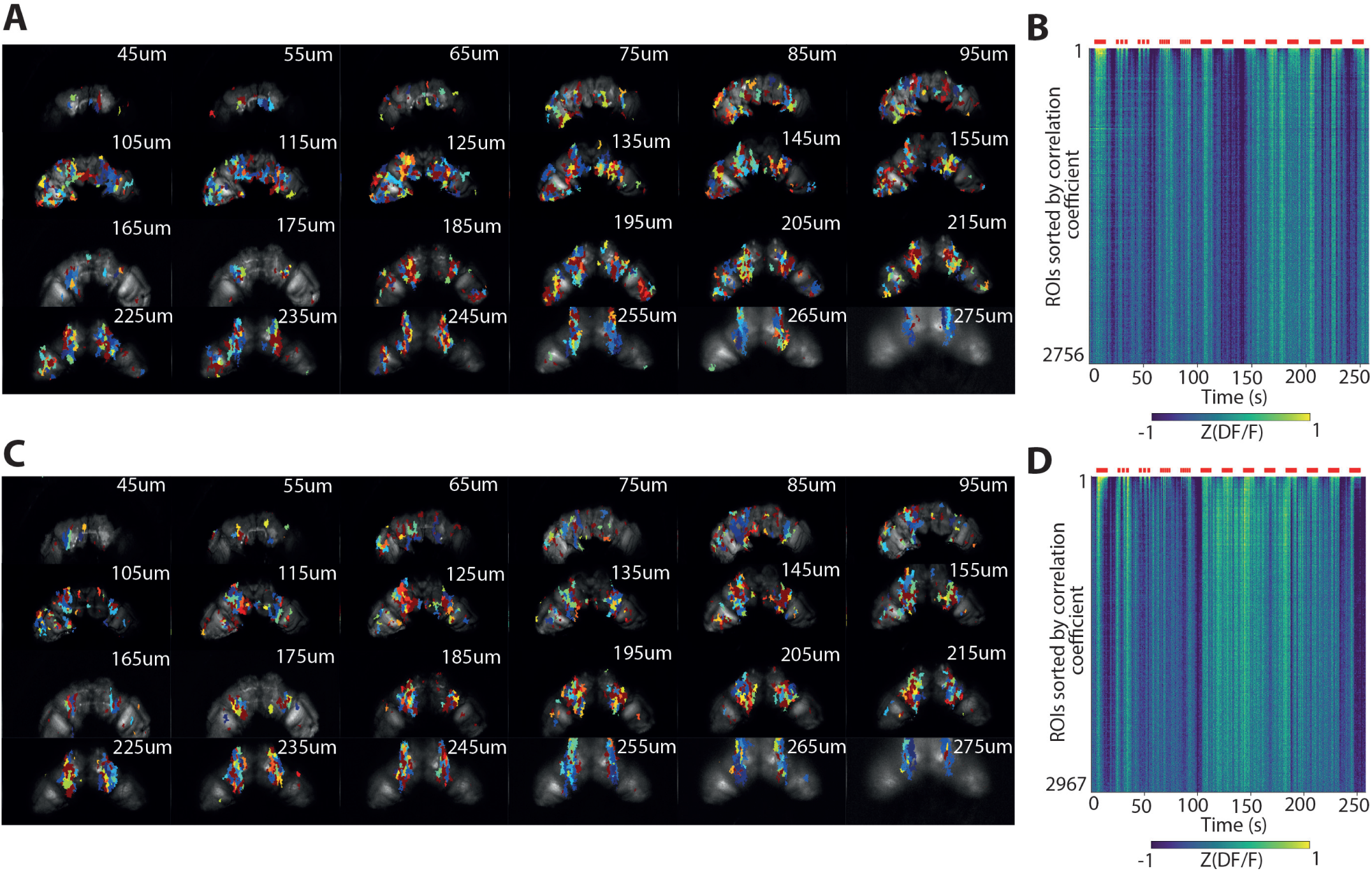
Auditory ROIs are widespread: **A)** Average intensity z-projection for each of the 24 planes for a representative brain extracted with LBM and overlaid with auditory ROIs. Number indicates depth in the sample. **B)** Heatmap of the activity of auditory ROIs sorted by correlation coefficient with the auditory stimulus (largest on top). **C-D)** Same as in A-B for a second representative brain.

**Figure S6.**
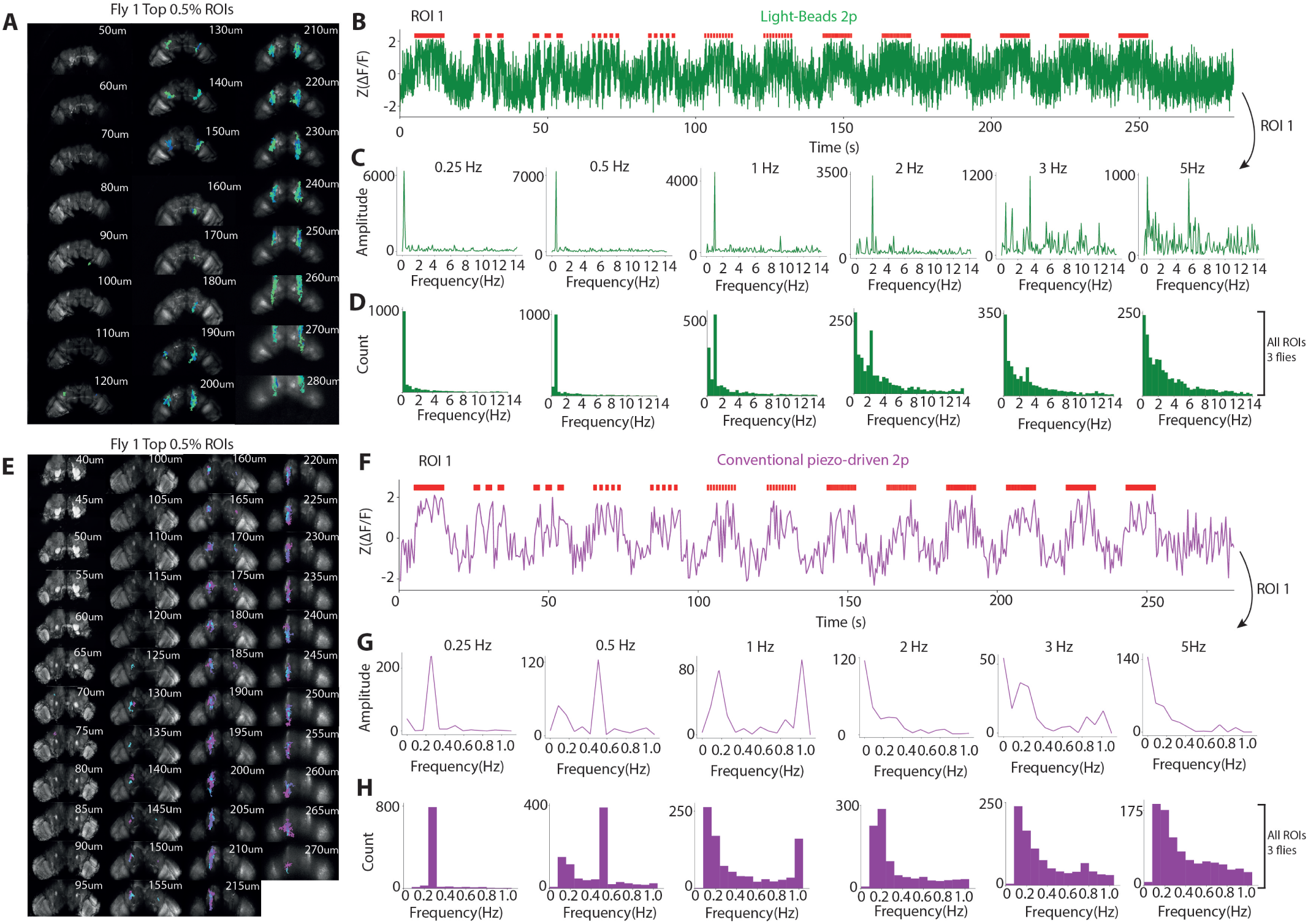
Single ROIs are time-locked to the stimulus. **A)** Average intensity z-projection for each of the 24 planes for a representative brain extracted with LBM and overlayed with the top 0.5% of ROIs with highest correlation coefficient. **B)** z-scored DF/F over time for a single ROI extracted with LBM. **C)** Fourier spectrum computed on each block at the indicated frequency. **D)** Distribution of Fourier peaks computed across all blocks of all ROIs across 3 animals. **E-H)**Same for conventional piezo-driven 2p.

**Figure S7.**
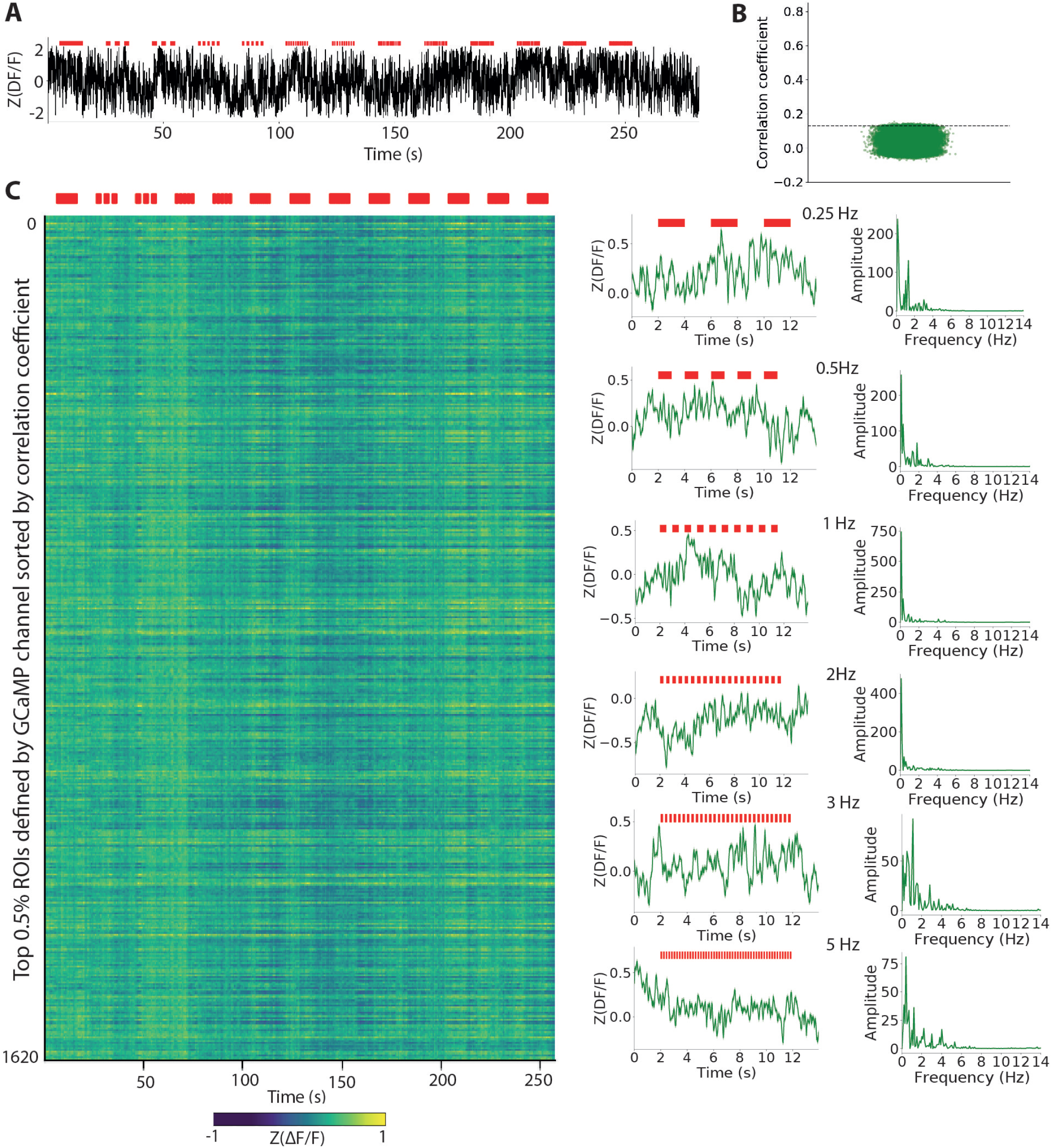
No stimulus-correlated ROIs are found in the red channel. Correlations with respect to the stimulus do not appear in the red channel. A) z-scored Delta F/F from a single ROI in the red channel. This ROI corresponds to the ROI with the highest correlation coefficient in the green channel. B) Top 0.5% of ROIs with the highest correlation to the convolved stimulus in the red channel. C) (Left) Activity of the top 0.5% of ROIs sorted by correlation coefficient (highest on top) extracted with LBM from the green channel and projected onto the red channel. (Middle) Z-scored mean activity of the top 0.5% ROIs during each stimulus block of the indicated frequency. Shaded area represents standard error of the mean (SEM). (Right) Fourier spectrum of the z-scored mean activity during each block.

**Figure S8.**
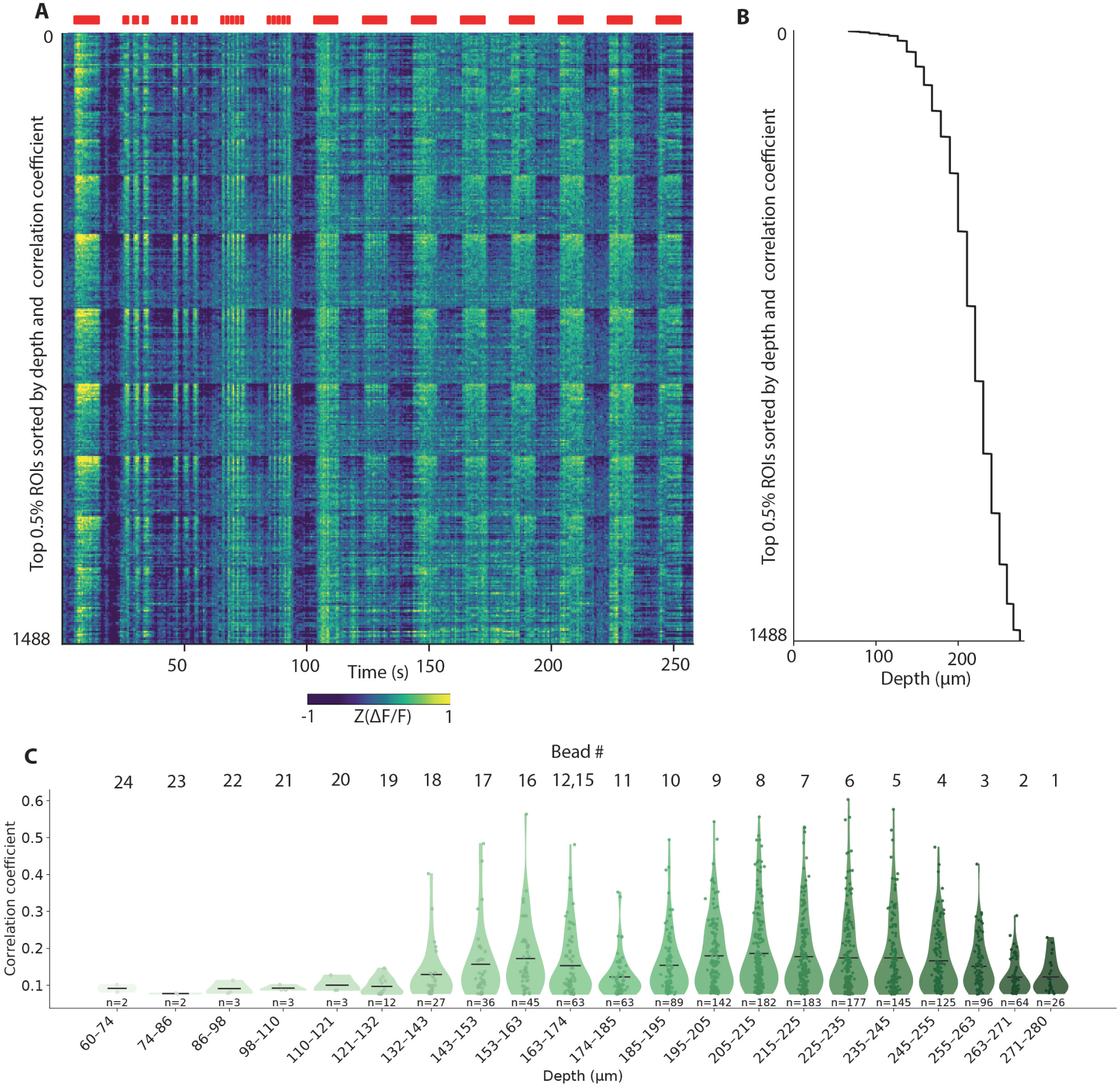
Auditory ROIs at different depths within the brain. A) Activity of the top 0.5% of ROIs extracted with LBM and sorted primarily by depth and secondarily by correlation coefficient (highest on top for each depth). B) Plot showing the depth of each ROI in A. C) Violin plot showing the correlation coefficient vs depth for each ROI in A.

**Figure S9.**
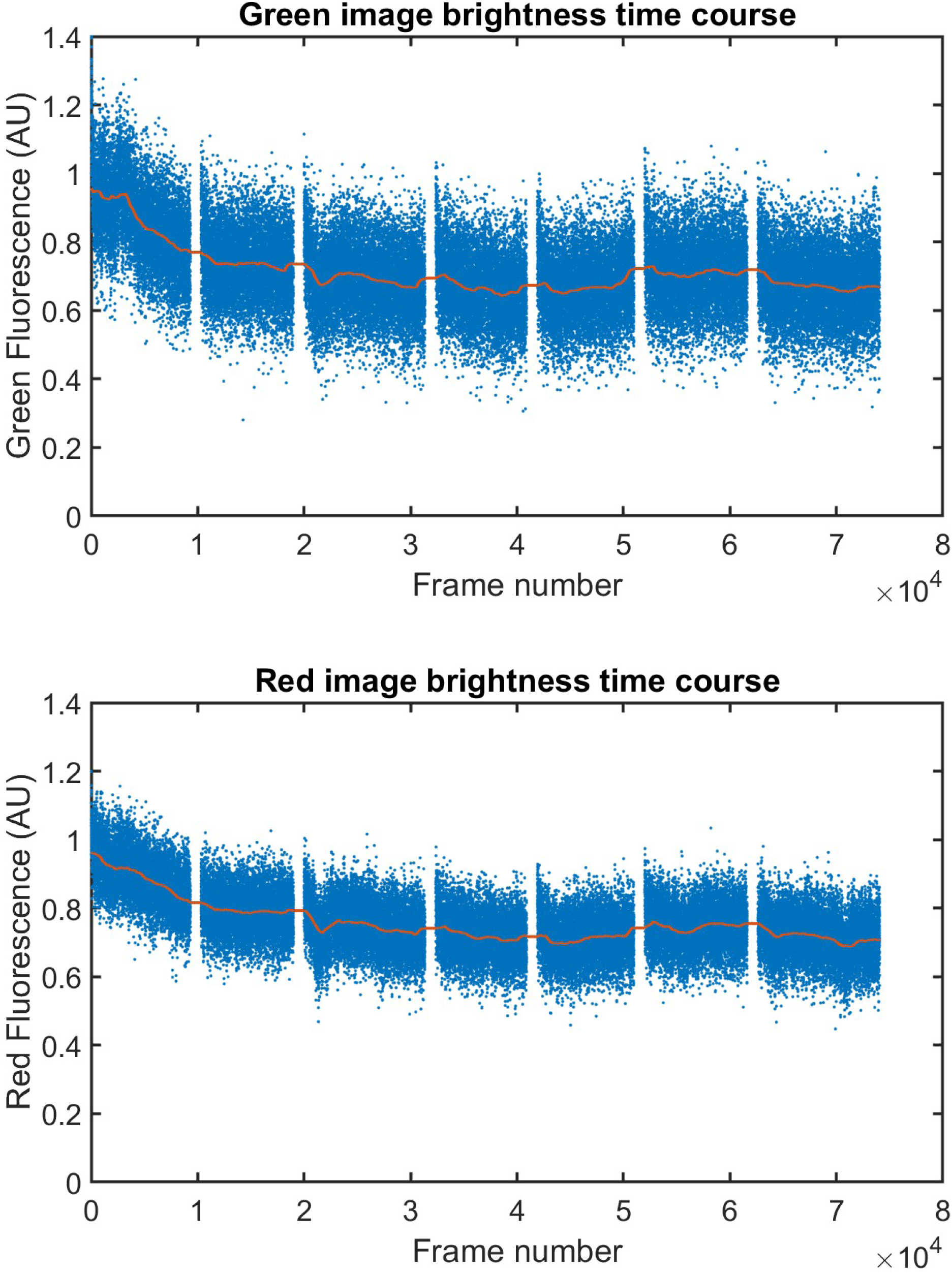
Assessment of photobleaching: Representative time course of the mean fluorescence in the central brain, in both the green and the red channels, here measured in plane 10. The time axis is expressed in frame number (acquired at 30Hz), and covers the duration of 7 trials (about 45min), more than 2x longer than in our experiments. Both green and red channels lose only about 25% of their average fluorescence signal.

**Figure S10.**
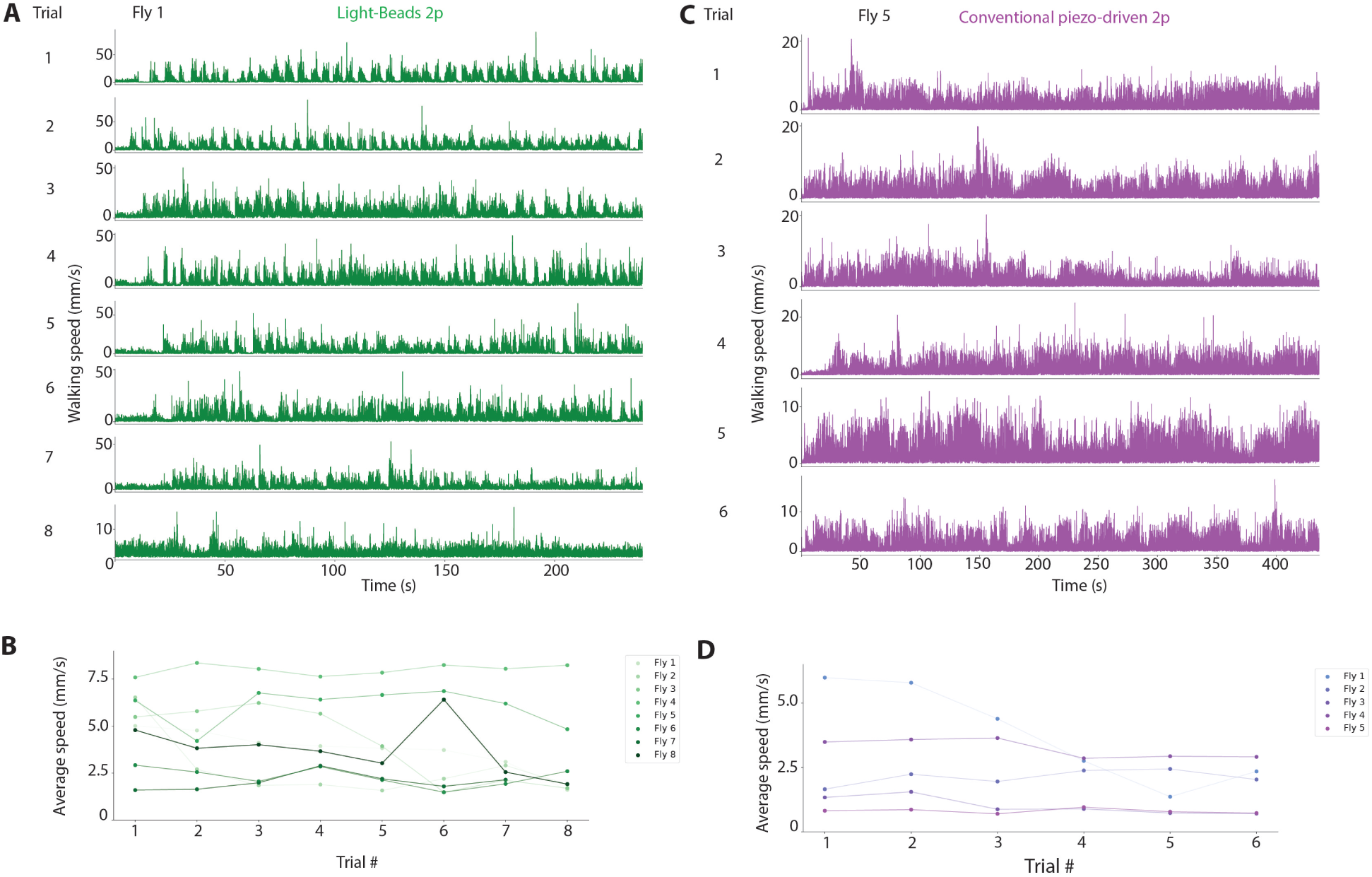
Animal health is stable across 40min recording. **A)** Plot of walking speed over time for 8 successive trials for a given fly recorded with LBM. **B)** Quantification of average speed across trials for 8 flies recorded with LBM. **C-D** Same for conventional piezo-driven 2p for 5 flies.

**Figure S11.**
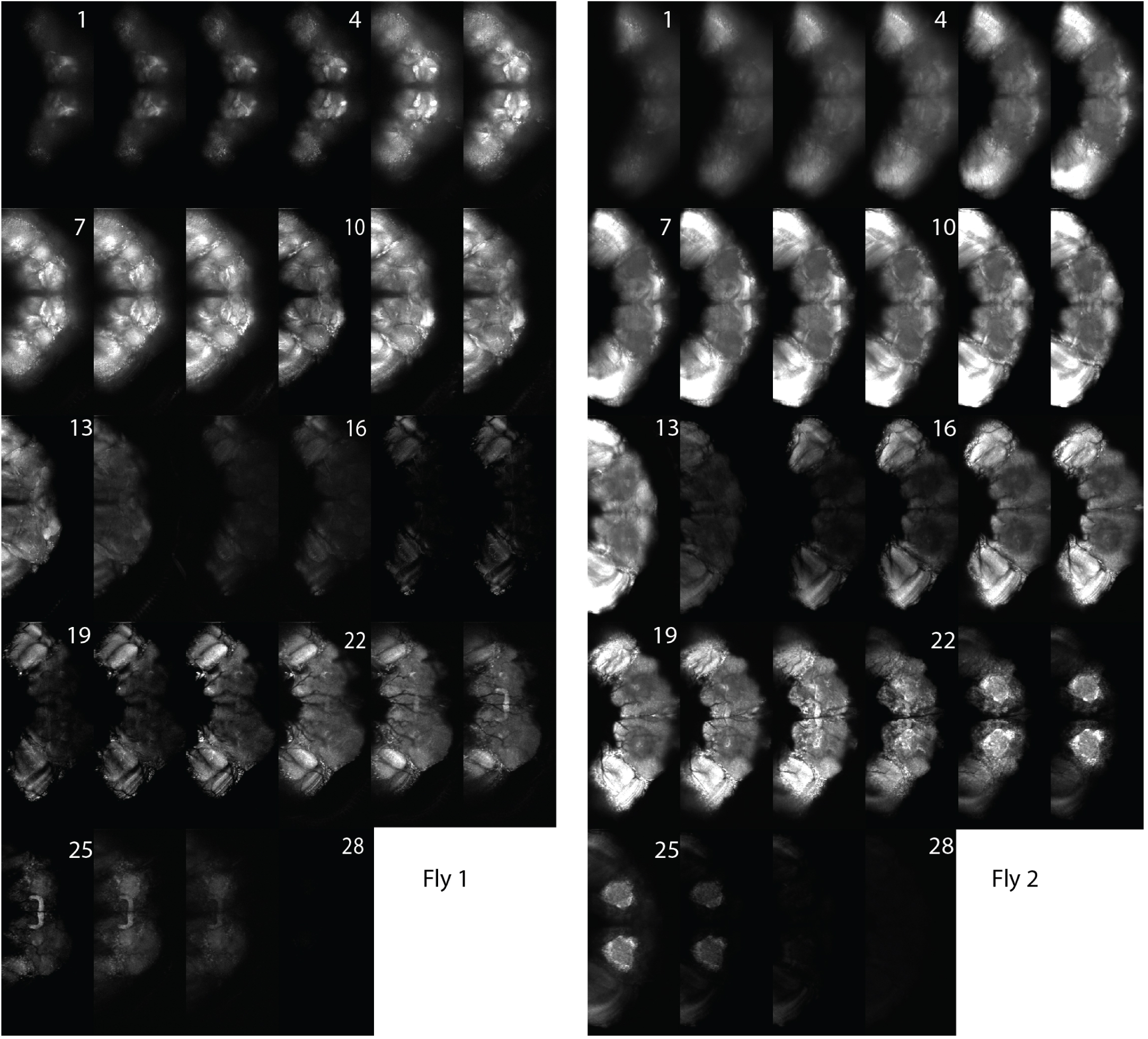
Planes brightness: The panels present the average images of each of the 28 planes in 2 separate experiments. All planes are displayed with the same contrast and brightness look-up table to allow for direct comparison of the intensities. Ideally, all planes would have the same intensity. However, multiple elements participate in determining the brightness of a given plane. The non-ideal value of the transmission of the beam splitter used in the cavity generates a gradient of intensity. However, the morphology and possibly the physiology of the fly brain generate variations as well. Channels 13, 14, 27 and 28 were not used in this study.

## Notes

### Competing Interest Statement

The authors have declared no competing interest.

### Summary of Updates

New Fig. 1: Optical layout, electronic and the logic of the signal collection; New Fig. S3 and S4: Images of sparse or pan-neuronal GCaMP lines; New Fig. S8: Stimuli-responsive ROIs when sorted by depth are found to primarily accumulate in deeper brain regions; New Fig. S9: Time-course of signal intensity across a long imaging session demonstrates low photobleaching levels; New Fig. S11: Signal intensity in the different planes.

